# Parabrachial oxytocin receptor-expressing neurons link social observation of distress to defensive behavior

**DOI:** 10.64898/2026.08.06.743398

**Authors:** Elena N. Judd, Jordan L. Pauli, Sayaka J. Kenmochi, Michael R. Bruchas, Richard D. Palmiter

## Abstract

The ability to detect and respond to threat signals in the environment, including those conveyed by the distress of a familiar social partner, is fundamental to survival and disrupted in a range of neuropsychiatric conditions. This study identifies oxytocin receptor (Oxtr)-expressing neurons within the lateral parabrachial nucleus (lPBN) of mice as a key node in the neural circuitry underlying threat-related and social behaviors. These Oxtr neurons are activated by aversive stimuli and by observing demonstrator mice in stressful situations, including foot shock or inflammatory pain. Chemogenetic inhibition of these neurons alters social proximity and pain contagion in observers without affecting general anxiety-like behavior. Inhibition also transiently suppresses non-social central sensitization. Direct activation of lPBN^Oxtr^ neurons with a selective Oxtr agonist is anxiogenic and results in increased tactile sensitivity. Together, these findings suggest that lPBN^Oxtr^ neurons are poised to integrate information about environmental threat, whether experienced directly or witnessed in a conspecific to coordinate appropriate defensive behavioral responses.

## INTRODUCTION

Oxytocin is a highly conserved hypothalamic neuropeptide with diverse neuromodulatory roles. Oxytocin receptor (Oxtr) signaling is widely appreciated for its role in parental and social pair-bonding behaviors^1–7^, but recent evidence suggests that its role within the central nervous system is complex and its impact on behavior depends both on social or environmental context and on the region of release^8^. Though oxytocin is often considered to impart positive affect and generally reduce pain-related behavior, it is also hypothesized to strengthen fear or escape responses to immediate threats^9–12^. Oxytocin signaling also regulates homeostatic processes such as food and water consumption^13–16^, thermoregulation^13,17,18^, and respiration^19^. Social or observational learning and emotional discrimination in many animal species require oxytocin receptor expression and is enhanced by oxytocin administration^20–25^.

Neurons that produce oxytocin largely reside within the supraoptic (SON) or periventricular hypothalamus (PVH)^26,27^. Magnocellular oxytocin neurons project mainly to the pituitary gland for hormonal release into the bloodstream while parvicellular oxytocin neurons project to a diverse array of central nervous system targets^28^. The oxytocin receptor is expressed broadly throughout the central and peripheral nervous system, but axonal mapping of oxytocin-producing neurons has shown more limited terminal fields^26^. For example, many cortical regions have extensive receptor expression but little evidence of nearby synaptic terminals. This has led to speculation of extra-synaptic transmission, including somato-dendritic or ventricular release, whereby oxytocin peptide is released in large quantities and acts on its receptor at slow timescales through diffusion^26,28,29^. In contrast, oxytocin positive terminals are relatively dense in the brainstem, including in the dorsal pons, but few studies have described a role for oxytocin receptor signaling in this region.

The lateral parabrachial nucleus (lPBN), a brain region in the dorsal pons, has access to sensory information of all modalities^30,31^ and is both necessary and sufficient for aversive responses to negative stimuli, as well as the cues that predict them^32,33^. lPBN activity, especially in *Calca*-expressing neurons, can induce peripheral pain sensitization in the absence of tissue damage^34,35^. lPBN neurons, including a subset of *Calca* neurons, express *Oxtr* and receive monosynaptic input from oxytocin-expressing neurons in the hypothalamus^14^. These PVH oxytocin inputs are glutamatergic and bath application of selective Oxtr agonists enhances spontaneous action potentials of lPBN^Oxtr^ neurons in slices. lPBN^Oxtr^ neurons modulate liquid consumption behavior but this effect may not be specific to oxytocinergic input^14,36^. The conditions that drive oxytocin release in this region or how oxytocin signaling in this area might influence behavior has not been described. Given the role of the lPBN in the affective processing of noxious stimuli^31,37^, we investigated the activity of lPBN^Oxtr^ neurons to aversive events and the observation of aversive events where the stressor (either foot shocks or inflammatory pain) was instead presented to a familiar social partner, the demonstrator.

Pioneering studies have established that brief interaction with a demonstrator undergoing visceral or inflammatory pain is sufficient to induce pain phenotypes in observer animals, mirroring though not fully recapitulating the sensitized state of the demonstrator^38–42^. Complementary work in the domain of observational fear conditioning has demonstrated that the social transmission of affective states extends beyond pain to encompass spontaneous and learned defensive responses more broadly. In this paradigm, an observer animal acquires freezing behavior upon witnessing a demonstrator receive aversive foot shocks, without any direct experience of the unconditioned stimulus itself^21,38,43–47^. These socially transferred phenotypes are modulated by familiarity, sex, and housing conditions, and is accompanied by anxiety-like behavior in the observer, consistent with a broad shift toward a hypervigilant or defensive behavioral state rather than a modality-specific nociceptive response^42,48–50^. Importantly, while observers reliably acquire freezing responses over repeated sessions, they often fail to form durable conditioned associations to discrete predictive cues to the same degree as directly shocked animals. This may reflect weaker plasticity or could imply that the neural mechanisms supporting observational fear learning diverge from those underlying standard associative conditioning^51^. Across both paradigms, observer behavioral responses are acquired during or shortly after social interaction with a distressed partner and do not require direct tissue damage or noxious stimulation, positioning them as ethologically critical examples of predictive threat-state adoption that may confer survival advantage in social species but become maladaptive when chronically or inappropriately engaged^52–54^. The anterior cingulate cortex, basolateral amygdala, nucleus accumbens, and their interconnections have been identified as critical nodes in the circuitry underlying both observational transfer of fear and pain^38,45,46,55–57^. Oxytocin signaling is also implicated in enhancing the fidelity and generalization of observationally acquired responses^21,22,24,58,59^. Whether subcortical regions with privileged access to ascending nociceptive and interoceptive signals, such as the lPBN, contribute to the initial neural encoding of observed threat that drives these behavioral outcomes has not previously been examined.

Here, we combine *in vivo* and *ex vivo* fluorescence recordings, targeted neural manipulations, region-specific drug delivery, and circuit-level anatomical tracing to interrogate the role of oxytocin signaling within the lPBN. We investigated a range of directly experienced and socially observed threats, revealing an unexpected contribution of this brainstem Otxr-expressing population to the neural mechanisms by which distress shapes defensive behavior. These findings suggest lPBN^Oxtr^ neurons are broadly responsive to threatening and aversive experience and reveal neural circuit mechanisms for how observed distress shapes defensive behavior in the witness.

## RESULTS

### lPBN^Oxtr^ neurons form a distinct population broadly recruited by aversive experience

To establish the molecular identity of lPBN^Oxtr^ neurons relative to previously characterized parabrachial subpopulations, we performed fluorescent in situ hybridization (FISH) with probes targeting *Oxtr*, *Slc17a6* (Vglut2), *Slc32a1* (Vgat), *Pdyn*, and *Calca*, (**Fig. 1a-d**).

**Figure 1.**
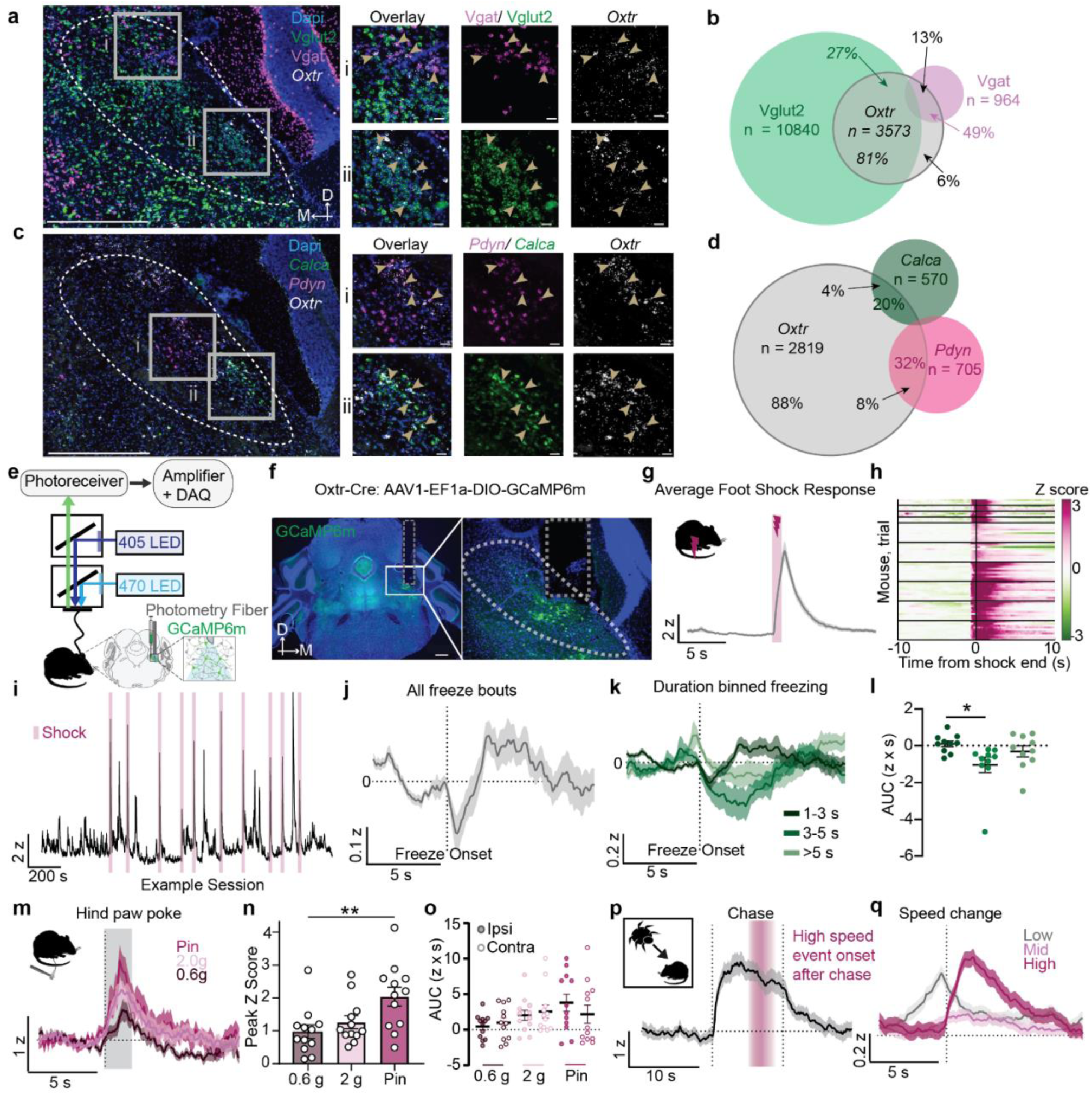
lPBN^Oxtr^ neurons represent an anatomically distributed population that are robustly activated by aversive stimuli and escape locomotion. **a.** Example fluorescent images from in situ hybridization co-labeling *Oxtr,* Vglut2 (*Slc16a6*), and Vgat (*Slc32a1*). Gold arrows depict cells with overlap. Scale bars are 500 μm (left) and 50 μm (right). **b.** Quantification of the cells (total number of cells = n) identified to be expressing each marker and the proportion of co-expression with each marker. **c, d.** Same as a and b but for in situ hybridization co-labeling *Oxtr, Calca*, and *Pdyn*. The total number of cells comes from 4-5 PBN sections across 3 (*Oxtr, Calca, Pdyn*) or 4 mice (*Oxtr*, Vgat, Vglut2). **e.** Schematic of fiber photometry recording with dual wavelength LED. **f.** Example GCaMP6m expression and fiber placement over lPBN in Oxtr-Cre mice. **g.** Peri-event averages of calcium fluorescence aligned to foot shock end (n =10). **h.** Heat map depicting each trial separated by individual. **i.** Z-scored trace from one example mouse over one foot shock recording session. 10 foot shocks were delivered with a variable ITI between 30 and 120 s with an average of 90s. **j.** Peri-event averages of calcium fluorescence aligned to freeze onset as detected by EzTrack. **k.** Same as H but analysis is binned by freeze bout length. **l.** Quantification of the difference in the AUC of the 5 seconds post freeze onset and the 5 s before freeze onset. **m.** Peri-event averages of calcium fluorescence aligned to the onset of hind paw stimulation lasting approximately 2 s (gray bar; n =12). **n.** Quantification of the peak Z-score of the data presented in m. **o.** Quantification of the AUC 5 s after paw stimulation onset; split by neural responses to ipsilateral and contralateral paw (relative to recording site). **p.** Peri-event averages of calcium fluorescence aligned to onset of chase periods (approximately 10 s, n =5). The red bar depicts the average time of high-speed escape onset after chase onset and the gradient reflects the range. **q.** Peri-event averages of calcium fluorescence aligned to speed change onset and binned by speed as detected by Ethovision. All fluorescence responses depicted as mean +/- SEM and z-scored over the full session.

Arrowheads in the high-magnification insets denote representative co-expressing cells across subregional sampling zones (**Fig. 1a**). Although the predominant neurotransmitter phenotype of lPBN neurons is glutamatergic, a subset of lPBN^Oxtr^ neurons co-expressed the vesicular GABA transporter *Slc32a1* (Vgat), indicating that a minority (∼13%) of this population is inhibitory (**Fig. 1b**). These data establish lPBN^Oxtr^ neurons as a dispersed, predominantly glutamatergic (∼81%) population that could respond to oxytocinergic input. *Pdyn*-expressing neurons, which mark dynorphin-producing cells of the dorsal lateral parabrachial nucleus, and *Calca*-expressing neurons, which label CGRP-producing cells of the external lateral parabrachial subdivision, define non-overlapping populations with distinct projection patterns and behavioral effects^60^. Quantification of co-expression revealed that the large majority of lPBN^Oxtr^ neurons (88%) do not co-express either *Pdyn* or *Calca*, representing a molecularly distinct population (**Fig. 1d**). Conversely, a portion of *Pdyn*- or *Calca*-expressing neurons co-expressed *Oxtr* (32% and 20%, respectively), indicating limited overlap with these well-characterized subpopulations. The connectivity and Oxtr-sensitive pharmacological properties of this population are characterized further in Figure 5; here we first focused on the activity profiles of lPBN^Oxtr^ neurons across a range of directly experienced threats.

Having established the molecular identity and neurotransmitter phenotype of lPBN^Oxtr^ neurons, we asked whether this population is functionally engaged during aversive events that the lPBN is known to process. To characterize the activity profiles of lPBN^Oxtr^ neurons during stressful events, we used adeno-associated viruses (AAVs) to express the genetically encoded calcium indicator GCaMP6m in lPBN of *Oxtr*^Cre^ mice and implanted fiber-optic cannulae dorsal to the lPBN^Oxtr^ for bulk fluorescence recordings via fiber photometry (**Fig. 1e**). Histological verification confirmed GCaMP6m expression and fiber tip placement and fluorescent labeling within the lPBN (**Fig. 1f, Extended Fig. 1**). This approach allowed us to monitor population-level calcium dynamics in lPBN^Oxtr^ neurons across a range of nociceptive and anxiogenic paradigms. We first examined neural responses to acute foot shock epochs, a well-established model of direct aversive stimulation. lPBN^Oxtr^ neurons exhibited robust, event-locked increases in GCaMP6m fluorescence following each shock delivery, as illustrated by the peri-event average trace (**Fig. 1g**) and the per-animal heatmap, which revealed consistent activation across individual subjects (**Fig. 1h**). An example trace aligned to repeated shock trials establishes the reliability and reproducibility of this response across multiple stimulations within the same animal with large fluorescence transients coinciding precisely with shock onset (**Fig. 1i**). These data demonstrate that lPBN^Oxtr^ neuron activity is recruited by acute noxious electrical stimulation.

In contrast to the robust activation observed during shock delivery, analysis of spontaneous behavioral state during the recording session revealed an opposing relationship between lPBN^Oxtr^ neuron activity and immobility. We measured freezing, the classical behavioral output of fear in rodents, by using machine-vision-assisted quantification (ezTrack)^61^. We found that tethered animals displayed an increase in freezing levels during the last five minutes of the foot-shock session compared to the first five minutes where no shock was given (**Extended Fig. 1**). Calcium sensor fluorescence intensity decreased upon onset of freezing bouts detected by this method (**Fig. 1j**) and further analysis indicates that this inhibition negatively scales with freezing bout duration, though long freezing bouts (>5 s) were comparatively rare (**Figure 1k-l**).

We then assessed whether lPBN^Oxtr^ neurons also respond to the intensity of mechanical stimuli applied to the plantar surface of the hind paw using von Frey filaments of increasing force and a pin prick. GCaMP6m fluorescence responses scaled positively with stimulus intensity, with the highest forces (pin and 2.0-g filaments) eliciting the largest peak calcium transient compared to lower-force stimuli (0.6-g filaments; **Fig. 1m**). Quantification of peak fluorescence amplitude across filament weights confirmed a significant relationship between stimulus intensity and neural activation (**Fig. 1n**). We found that the scaling of neural activity with force was unrelated to the laterality of hind paw stimulation relative to the recorded hemisphere (**Fig. 1o**). Taken together, these results indicate that lPBN^Oxtr^ neuron activity reflects the magnitude of peripheral tactile stimulation.

To determine whether lPBN^Oxtr^ neurons respond to ethologically relevant threat in the absence of physical harm, we examined calcium dynamics during a simulated predator paradigm in which a large, robotic six-armed bug modeled predator approach, reliably eliciting escape behavior in mice. lPBN^Oxtr^ neurons showed significant increases in GCaMP6m fluorescence as the toy robot approached (**Fig. 1p**). Speed-stratified averaging revealed population activity was markedly higher during high-speed locomotion events (>10 cm/s) compared to mid (1.5–10 cm/s) or low-speed epochs (<1.5 cm/s; **Fig. 1q**). All high-speed events (>10 cm/s) were associated with increased calcium fluorescence at their onset, but those that arose during predator-chase periods exhibited higher intensity than those that occurred while the ‘predator’ was stationary (**Extended Fig.1**). Our finding that lPBN^Oxtr^ neurons are activated both during the anticipatory period preceding escape and during high-speed flight itself suggests that these neurons integrate a composite signal reflecting both threat imminence and the subsequent motor response.

We also investigated whether lPBN^Oxtr^ neuron activity was modulated during an anxiogenic paradigm that does not involve physical perturbation. Animals were placed in an elevated plus maze (EPM), which exploits the innate conflict between exploration and avoidance of exposed, open spaces. Average GCaMP6m traces revealed that lPBN^Oxtr^ neurons exhibited differential activity during open versus closed arm occupancy, with elevated fluorescence observed during closed arm entries relative to the open arm entries (**Extended Fig.1**). Quantification of time spent in each arm showed a robust preference for the closed arm over the center area or open arms (**Extended Fig.1**). We hypothesized that this effect may be due to the speed at which the animals entered the different compartments of the behavioral apparatus. Quantification of speed at zone entry showed that animals enter closed arm zones at higher speed than open arm zones (**Extended Fig.1**). Like in the predator chase experiment, the EPM experiment showed a positive relationship between locomotion velocity and GCaMP6m signal magnitude. Conversely, we found that a 10-s tail suspension, an innately stressful but locomotion-independent test, elevates lPBN^Oxtr^ neuron activity (**Extended Fig.1**). These findings suggest that lPBN^Oxtr^ neuron activity is not exclusively driven by nociceptive input. While a component of this signal co-varies with locomotion velocity, these neurons can also be recruited by anxiogenic stimuli independent of movement. The convergence of nociceptive, anxiogenic, and threat-related signals onto this population, combined with the tight coupling of their activity to locomotion velocity, suggests that lPBN^Oxtr^ neurons occupy a functionally significant node at the intersection of sensory, affective, and motor systems, poised to integrate information about environmental threat and coordinate appropriate behavioral responses.

### Direct and observed inflammatory pain recruit lPBN^Oxtr^ neurons

lPBN^Oxtr^ neurons are broadly responsive to nociceptive and threat-related stimuli and since oxytocin is hypothesized to play important roles in social and observational contexts, we asked whether this population is similarly engaged when such stimuli are experienced vicariously through a social partner. We used a model of socially induced tactile sensitivity in which mice interacting with a familiar, same-sex partner undergoing inflammatory pain induced by subplantar injection of Complete Freund’s Adjuvant (CFA) exhibit tactile sensitivity that mimics, but is less severe than, that of their pained partner^38,39^ (**Fig. 2a**). Demonstrator animals receiving subplantar CFA exhibited marked and prolonged ipsilateral tactile sensitivity. A one-hour interaction with a stressed partner was sufficient to induce moderate bilateral allodynia in observers when assessed on the same day, an effect absent in pain-naïve control dyads where neither animal received CFA (**Fig. 2b, c**). When social interaction was limited to one hour and animals were subsequently cross housed with non-stressed partners, observer paw withdrawal thresholds (PWTs) returned to baseline. The one-hour social interaction was repeated four days post-CFA injection, at which point observers again exhibited mechanical sensitivity upon reunion with a pained conspecific (**Fig. 2b, c**). To characterize the activity profiles of these neurons during social observation itself using paired fiber photometry recordings of GCaMP6m fluorescence in lPBN^Oxtr^ neurons during the social interaction period following subplantar CFA injection to the demonstrator animal (**Fig. 2d**). Manual scoring of the interaction period for social investigation, allogrooming, and individual grooming bouts revealed an increase in observer-driven investigation and demonstrator grooming after subplantar injection (**Fig. 2 e, f**). Allogrooming was detected in some but not all pairs. The low number of instances of each behavioral class, combined with their frequent co-occurrence with reduced locomotion, contributed to high signal variability and precluded reliable event-aligned analysis of calcium dynamics during these epochs. Peri-event averages aligned to von Frey filament application at moderate (0.6 g) and high (2.0 g) forces in this context revealed time-locked responses comparable to those described in **Figure 1**, though no difference in neural response amplitude was detected as a function of pain induction or pain observation state (**Fig. 2 g-i).** We were unable to find evidence of enhanced neural responses to filament stimulation of the hind paw even when directly comparing the injured and uninjured paw in demonstrator mice. To quantify enhanced neural activity following CFA administration in this assay, we detected calcium transients exceeding the median absolute deviation of the fluorescence signal across the session (60) and assessed both their amplitude, measured as area under the curve (AUC), and their frequency in a 30-s rolling window across the session (**Fig. 2j, k**). CFA injection in the demonstrator enhanced both the amplitude and frequency of calcium transients in demonstrator animals, while observers exhibited a selective increase in transient amplitudes without a corresponding change in frequency (**Fig. 2k**).

**Figure 2.**
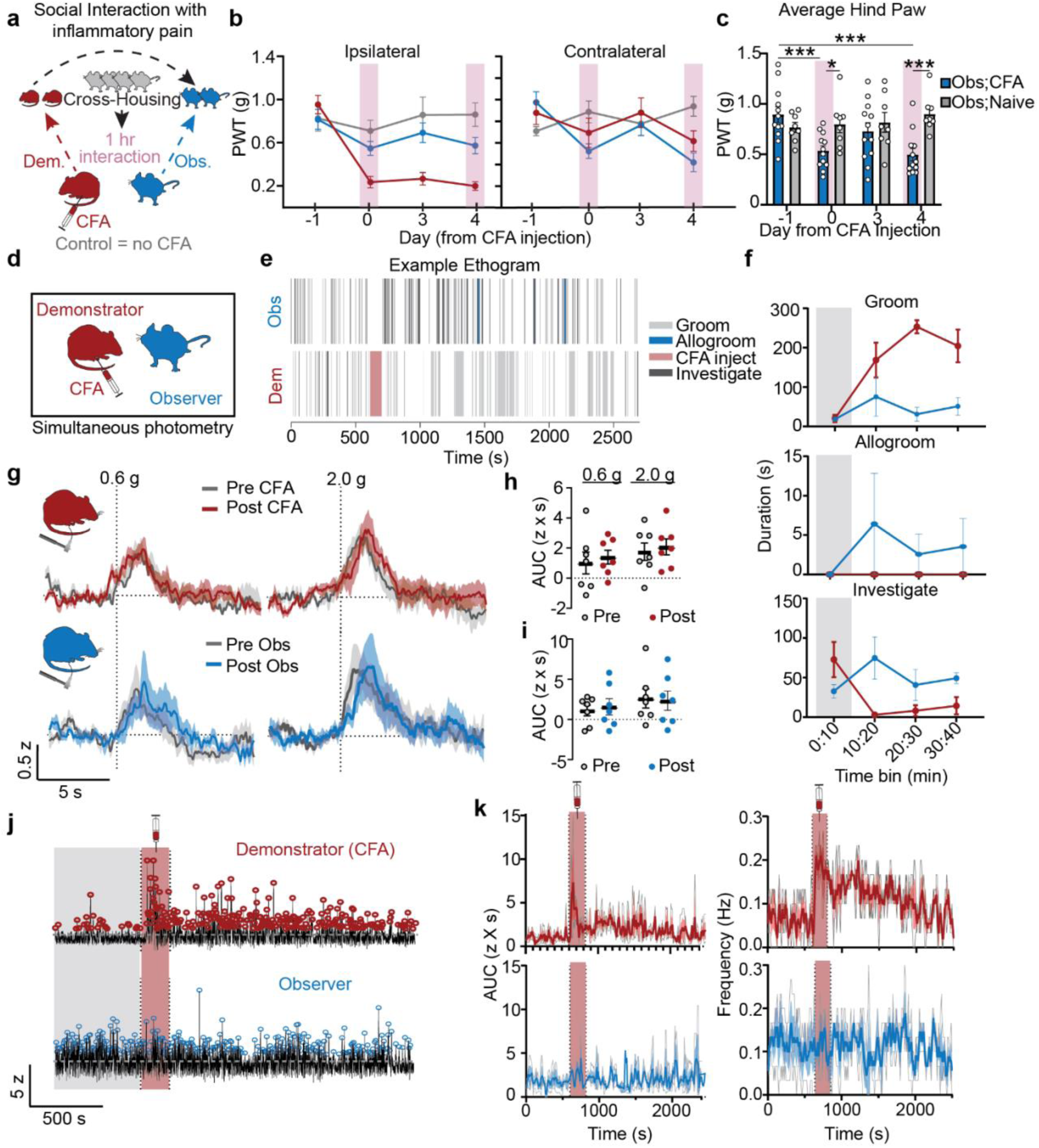
lPBN^Oxtr^ neurons are activated by observing demonstrator mice with inflammatory pain. **a.** Schematic depicting the cross-housing and social interaction protocol as detailed in Rein et al., 2022. **b.** PWT from the ipsilateral and contralateral paw that received CFA in demonstrator mice. Social interaction periods (pink) occurred on days 0 and 4. Neither mouse in the control pair received CFA. **c.** Quantification of the average of the hind paw PWT of observer mice only. Two-way ANOVA. **d.** Schematic of simultaneous fiber photometry recordings during social observation of inflammatory pain. **e.** Example ethogram from manually scored behavior during the social interaction recording window. **f.** Quantification of the time each animal in the dyad spent performing manually scored behaviors. Grey bars here denote baseline period before CFA was administered to the demonstrator. **g.** Peri-event averages of demonstrator calcium fluorescence aligned to the onset of hind paw stimulation with 0.6 g (left) and 2.0 g (right) von Frey filaments. Grey trace is before CFA, red is after. Below shows the same data is from CFA-observer mice (blue, n =7). **h.** Area under the curve (0-5 s) post hind paw stimulation for each filament weight, separated by trials acquired 24 hours before or after CFA injection. **i.** Same as h for CFA observer mice. **j.** Calcium transient analysis using peak detection over hour long recordings in demonstrator and observer mice. **k.** Quantification of the AUC of each detected peak (left) or peak frequency (right). Data calculated over a 30 s rolling window.

### lPBN^Oxtr^ neurons are activated by observed foot shock

To leverage a paradigm with a more structured, trial-based design better suited to photometry recordings, we turned to an observational fear conditioning model. A standard fear conditioning chamber was divided with a perforated, clear plexiglass barrier such that the demonstrator received ten 0.5-mA foot shocks while the observer remained on a non-electrified surface (**Fig. 3a**). Each foot shock was predicted by and co-terminated with a 10-s auditory tone serving as a conditioned stimulus (CS+). Both demonstrator and observer animals exhibited elevated freezing to the CS+ relative to pre-tone baseline, and observer freezing remained elevated throughout the first thirty seconds of the inter-trial interval (**Fig. 3b**). Calcium activity in observer lPBN^Oxtr^ neurons was significantly elevated during and following demonstrator foot shock delivery, and this elevation scaled with shock duration (**Fig. 3c**). This response was absent when both animals were pre-exposed to the tone without accompanying shock delivery, confirming that the neural response reflects processing of the aversive event rather than the auditory cue alone (**Fig. 3**, tone-alone condition). Significant increases in observer calcium activity during the CS+ period were not observed until a second conditioning session, consistent with behavioral experiments in which observer freezing to the CS+ did not exceed that of shock-naïve control observers until repeated exposure (**Extended Fig. 2**). In a subsequent test phase following two conditioning sessions, observers did not exhibit a significant neural response to the CS+ when assessed in isolation, whereas demonstrator animals did, suggesting that observers do not form lasting conditioned neural associations in this paradigm, consistent with the behavioral data (**Extended Fig. 2, 3**).

**Figure 3.**
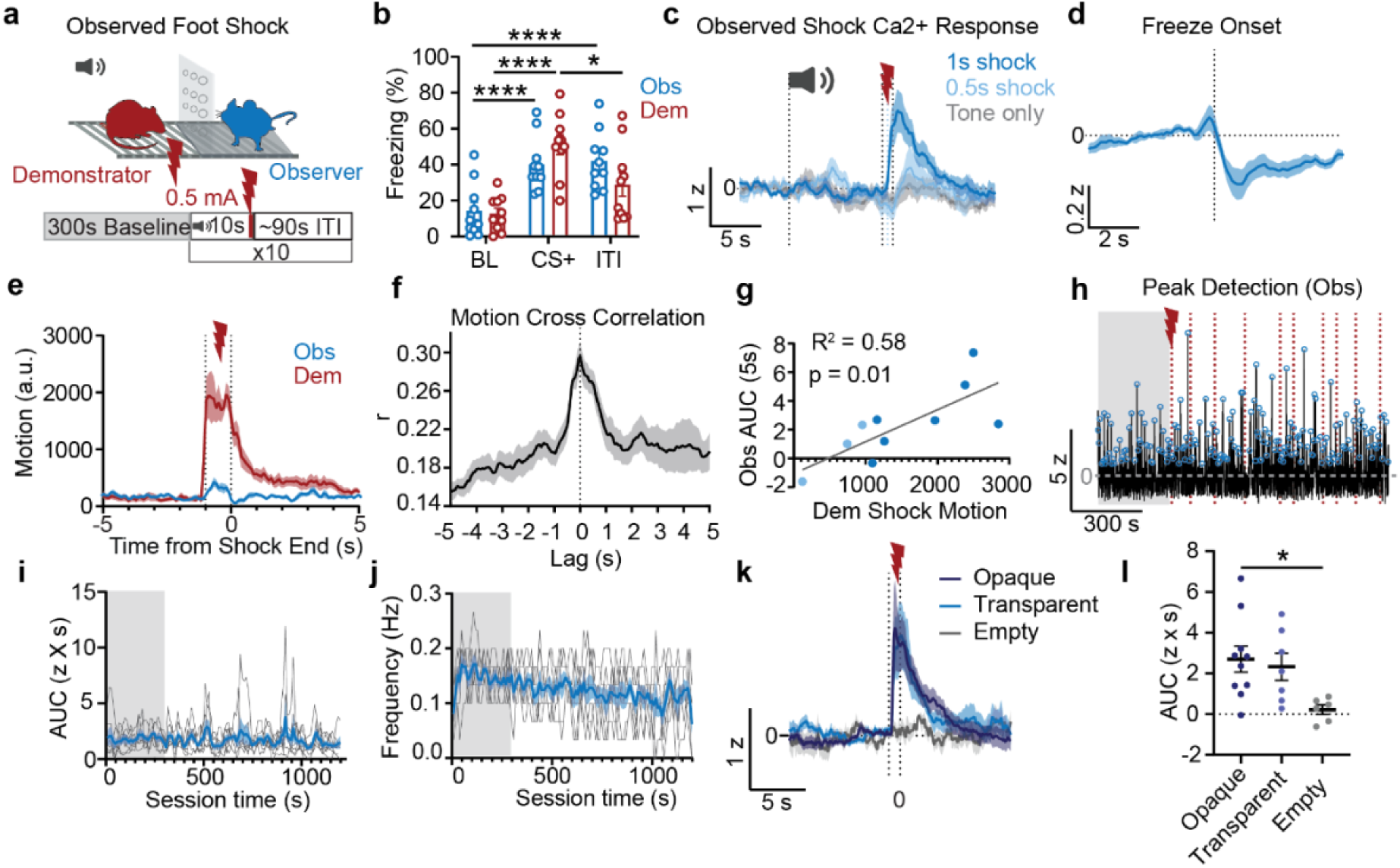
lPBN^Oxtr^ neurons are activated by observing foot shocks delivered to familiar partners. **a.** Schematic of observed foot shock paradigm. 10 s auditory cue serves as CS+. **b.** Quantification of freezing data extracted from ezTrack. **c.** Peri-event averages of observer calcium fluorescence aligned to the end of demonstrator foot shock delivery. Tone alone condition contained 4 CS+ presentations with no shock delivery (n =10), 1 s shock (n = 7) and 0.5 s shock (n= 3) contained 10 CS+ and shock trials each. All conditions had both mice present in the chamber. **d.** Peri-event averages of observer calcium fluorescence aligned to the onset of freezing bouts in observers (n = 10). **e.** Peri-event averages of motion calculated with ezTrack for all demonstrator and observer mice. **f.** Cross-correlation of the demonstrator and observer motion values shown in e. **g.** Pearson’s correlation of demonstrator motion response during the shock period with the observer AUC (0-5 s). **h.** Observer calcium transient analysis using peak detection over demonstrator foot shock session. **i.** Quantification of the AUC of each detected peak (left). Data calculated over a 30 s rolling window. **j.** Same as I but peak frequency plotted instead. Grey lines are individual mice, and blue is the group average +/- SEM. **k.** Peri-event averages of observer calcium fluorescence aligned to the end of un-cued demonstrator 1 s foot shock delivery. Separate sessions of 10 trials were completed with either an opaque or transparent divider as well as an additional control session where foot shocks were delivered but no demonstrator mouse was present on the shock grid (empty). **l.** Quantification of the AUC (0 to 5 s) in each condition shown in k.

As seen in the direct shock experiments of **Figure 1**, lPBN^Oxtr^ neurons in observers showed suppression of calcium activity at the onset of freezing (**Fig. 3d**). To determine the extent to which elevated neural activity during shock periods reflected coinciding movement, we quantified motion in demonstrator-observer pairs over matched tone and shock epochs using a pixel-change metric (ezTrack) (59). Both members of each pair showed elevated motion during the shock period, though the amplitude of motion was substantially lower in observers than demonstrators (**Fig. 3e**). Pairwise cross-correlation of demonstrator and observer motion traces, averaged across all recorded pairs (n = 10), revealed tight coupling between the motion of the two animals with a small positive lag (**Fig. 3f**), indicating that demonstrator movement slightly preceded that of the observer. This may be consistent with a demonstrator-induced startle response in observers. Regression analysis further revealed a significant positive correlation between demonstrator motion amplitude during the shock period and observer lPBN^Oxtr^ activity in the five seconds following shock onset (r² = 0.58), indicating that the magnitude of the behavioral response in the demonstrator is a meaningful predictor of the neural response recruited in the observer (**Fig. 3g**). When observer responses were stratified by sex, male observers exhibited larger neural responses than females, though this was accompanied by correspondingly larger demonstrator shock responses in male pairs (**Extended Fig. 2**).

Peak detection analysis of lPBN^Oxtr^ activity in shock observers, performed using the same approach applied to CFA observers, revealed a preferential increase in calcium transient amplitude rather than a change in frequency of events (**Fig. 3h-j**) suggesting that enhanced transient amplitude may be a conserved feature of lPBN^Oxtr^ engagement during observation of externally experienced threats, regardless of the modality or source of the stressor.

Finally, to assess the social signals that are necessary for the observer neural response, we used un-cued 0.5-mA, demonstrator foot shocks. Neural responses in observers were comparable in magnitude to those observed in the cued paradigm, confirming that lPBN^Oxtr^ activity is driven by the shock event itself rather than by anticipatory cue processing **(Fig. 3k).** The observer calcium response to demonstrator foot shock was not primarily driven by visual input as we saw similarly enhanced GCaMP fluorescence when observers were behind an opaque rather than transparent divider (**Fig. 3k, l**). Other social cues, such as vocalizations, pheromones, or vibrations/ movement of the testing chamber may instead drive the observer response. Importantly, there was no appreciable neural response to shock delivery if the demonstrator side was empty (**Fig. 3l**). This confirms that the observer response is driven by shock-induced changes in the social partner rather than a sensory experience of the observer itself. Taken together, these data indicate that social observation of aversive events engages lPBN^Oxtr^ neurons in a manner that reflects both the severity and behavioral salience of the observed stressor. This positions lPBN^Oxtr^ neurons as a brainstem substrate through which witnessed threat may promote hypervigilant and nocifensive states in observers.

### Chemogenetic inhibition of lPBN^Oxtr^ neurons prevents tactile hypersensitivity in central pain sensitization

Social interaction with a pained partner elicits anxiety and pain-like behavior in mice, and lPBN^Oxtr^ neuron activity appears elevated by experienced and witnessed distress, therefore we asked whether lPBN^Oxtr^ neurons are required for the hypervigilant and nocifensive phenotypes in observers of inflammatory pain. To address this, *Oxtr^Cre^* mice were injected bilaterally in the lPBN with either an AAV carrying a Cre-dependent inhibitory hM4Di-mCherry (hM4Di) or a control (mCherry) construct and allowed to recover prior to behavioral testing (**Fig. 4a**). Histological verification confirmed expression of hM4Di-mCherry within the lPBN (**Fig. 4a, left**). Animals remained paired in their designated demonstrator-observer roles for the duration of the experimental timeline. The initial one-hour social interaction period following subplantar CFA injection to the demonstrator mouse captured the acquisition phase of observer hypersensitivity, while all subsequent days of co-housing provided ongoing exposure to the pained demonstrator. Mechanical sensitivity was assessed via von Frey-filament application before and after CFA administration to the demonstrator, and anxiety-like behavior was measured in the EPM on the same day as the social interaction period (**Fig. 4b**). The first and last ten minutes of each social-interaction period were video recorded and analyzed with proximity defined as less than 5 cm between the center points of the two animals.

**Figure 4.**
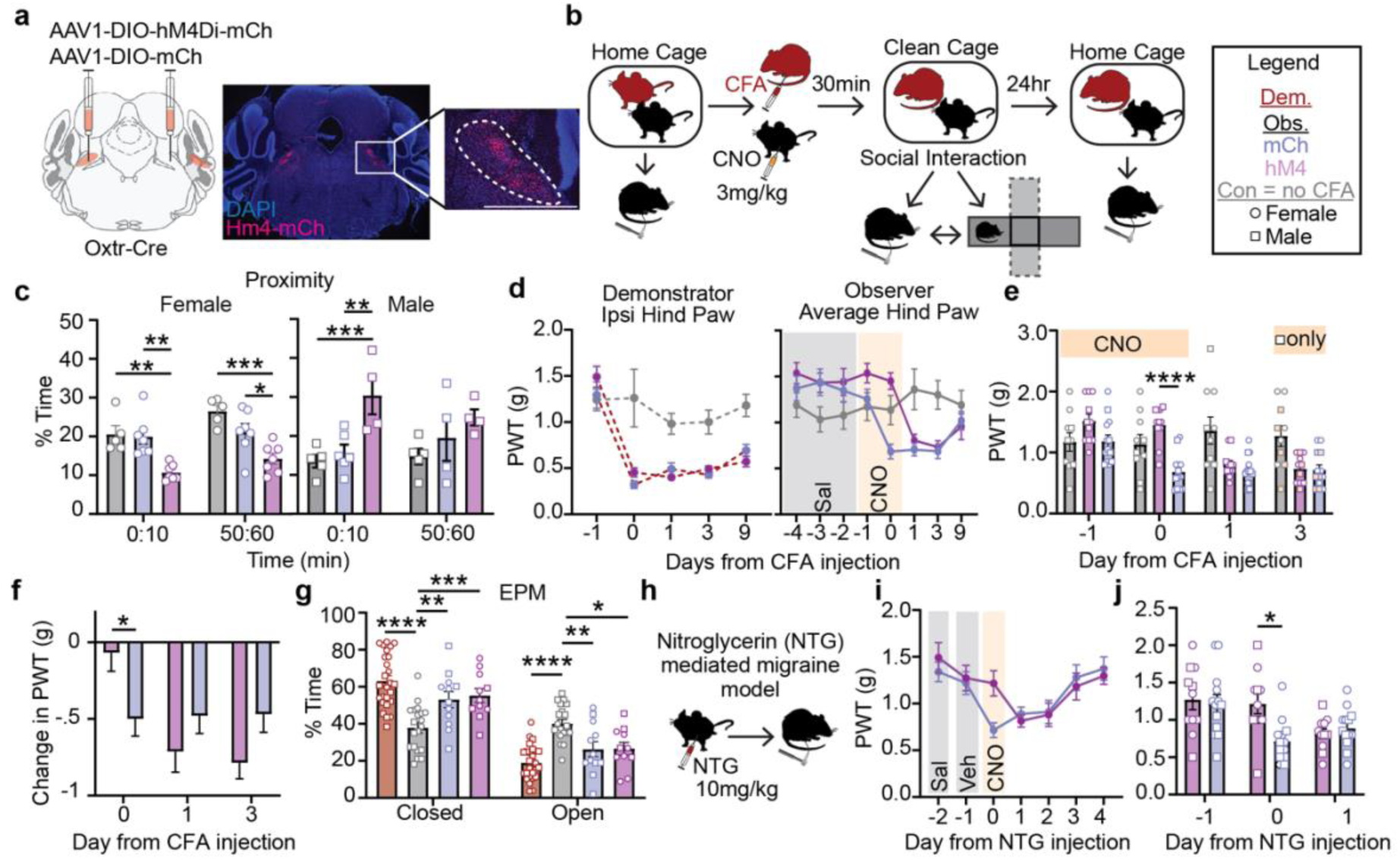
Chemogenetic inhibition of lPBN^Oxtr^ neurons reduces mechanical sensitivity in social and non-social paradigms of central sensitization. **a.** Schematic of bilateral virus injections and example fluorescence in lPBN of hM4-mCh. **b.** Schematic of home-cage paired dyads with CFA and CNO-administration. **c.** Time in proximity in the first 10 and last 10 minutes of the 1-hour social interaction period; females on the left, males on the right. **d.** Ipsilateral PWT of demonstrator mice (left) and average hind PWT of observer mice plotted as a function of day from demonstrator CFA injection (middle). **e.** Quantification of observer average PWTs -1 to 3 days from CFA injection. Two-way ANOVA. **f.** Quantification of the change in PWT from days 0, 1, or 3, relative to -1 for hM4 and mCh observers. Two-way ANOVA. **g.** Time in the open and closed zones in the EPM. Two-way ANOVA. **h.** Schematic of NTG-induced migraine model. **i.** Line graph of the average PWTs in mCherry and hM4 mice plotted as a function of days from NTG injection (I.P.). **j.** Quantification of data in i showing individual values. Control mice (grey) were treated the same way but neither individual in the dyad received CFA. Circle = female, square = male.

Chemogenetic inhibition of lPBN^Oxtr^ neurons via Clozapine-N-Oxide (CNO) administration during the social interaction phase disrupted affiliative behavior in a sex-specific manner. Inhibition reduced proximity in females (**Fig. 4c, left**) and increased proximity in males (**Fig.4c, right**), with effects most pronounced during the first ten minutes of the one-hour interaction period. At baseline, male dyads generally spent less time in proximity than females, reflecting possible sex differences in affiliative social behavior. No significant difference in proximity was observed between pain-naïve control pairs and mCherry control pairs given CFA in either sex. Despite the opposing effects of chemogenetic inhibition on proximity across sex, no sex difference in the mechanical sensitivity of observer or demonstrator mice was detected (**Extended Fig. 4**). Critically, chemogenetic inhibition of lPBN^Oxtr^ neurons prevented the acquisition of tactile sensitivity in observers (**Fig. 4d-f**). When hM4Di observers were assessed the following day after co-housing with the demonstrator in the absence of CNO, they displayed sensitivity comparable to mCherry controls. Observer sensitivity was largely resolved by nine days post-CFA injection, though demonstrator PWTs remained low, consistent with persistent sensitization (**Fig. 4d**). Inhibition of lPBN^Oxtr^ neurons in male animals after hypersensitivity was already established (Day 3 post-CFA) had no effect, indicating a specific role for this population in the acquisition, but not maintenance, of observer-induced hypersensitivity.

Although demonstrator and observer mice both showed elevated anxiety in the EPM relative to pain-naïve controls, chemogenetic inhibition of lPBN^Oxtr^ neurons did not significantly alter anxiety levels in this assay (**Fig. 4g**), suggesting that the contribution of this population to observer hypersensitivity may be dissociable from its effects on general anxiety-like behavior. Male animals showed modestly lower anxiety levels than females across groups, though overall trends were consistent across sexes. No differences in anxiety-like or nocifensive behaviors were observed when groups were separated by the timing of each assay relative to the initial social interaction (**Extended Fig. 4**).

To evaluate the specificity of lPBN^Oxtr^ neuron involvement in the development of socially transferred hypersensitivity, we also examined a non-social model of central sensitization. We turned to nitroglycerin (NTG) administration, an established rodent model of migraine that produces broad tactile sensitivity in both the orofacial region and the hind paws^35,62,63^ (**Fig. 4h**). Chemogenetic inhibition of lPBN^Oxtr^ neurons transiently suppressed the development of NTG-induced hypersensitivity, paralleling the effect observed in the social observation paradigm (**Fig. 4i, j**). Taken together, these findings suggest that lPBN^Oxtr^ neurons contribute to the acquisition of mechanical sensitivity across multiple models of central sensitization that occur in the absence of direct peripheral tissue damage.

### Oxytocin signaling within the lPBN is excitatory and promotes nocifensive behaviors

To directly establish whether lPBN^Oxtr^ neurons are functionally excited by oxytocin and to characterize the anatomical circuit this signaling may engage, we turned to *ex vivo* slice imaging, localized pharmacology, and circuit-level anatomical approaches. To corroborate evidence of functional excitation in lPBN^Oxtr^ neurons by oxytocin, we used two-photon, *ex vivo* slice imaging. AAVs delivering GCaMP8m were injected bilaterally into the lPBN of *Oxtr^Cre^* mice, enabling calcium imaging in acute brain slices (**Fig. 5a**). A micropipette was positioned in the vicinity of GCaMP-positive cells and used to deliver 200-ms pressure puffs of the selective oxytocin receptor agonist TGOT (Thr4,Gly7-oxytocin; 100 µM) while fluorescence was continuously monitored under two-photon illumination (**Fig. 5b**). A representative TGOT-evoked calcium response is shown in **Fig. 5c**. Puff application of artificial cerebrospinal fluid (aCSF) vehicle in place of TGOT did not produce comparable activation, establishing that the observed responses reflect specific receptor-mediated signaling rather than a mechanical tissue deflection artifact (**Fig. 5d, e**). As a positive control for slice health and GCaMP dynamics, recorded cells were also imaged during bath application of AMPA, a glutamate receptor agonist, which produced robust increases in fluorescence (**Extended Fig. 5**).

**Figure 5.**
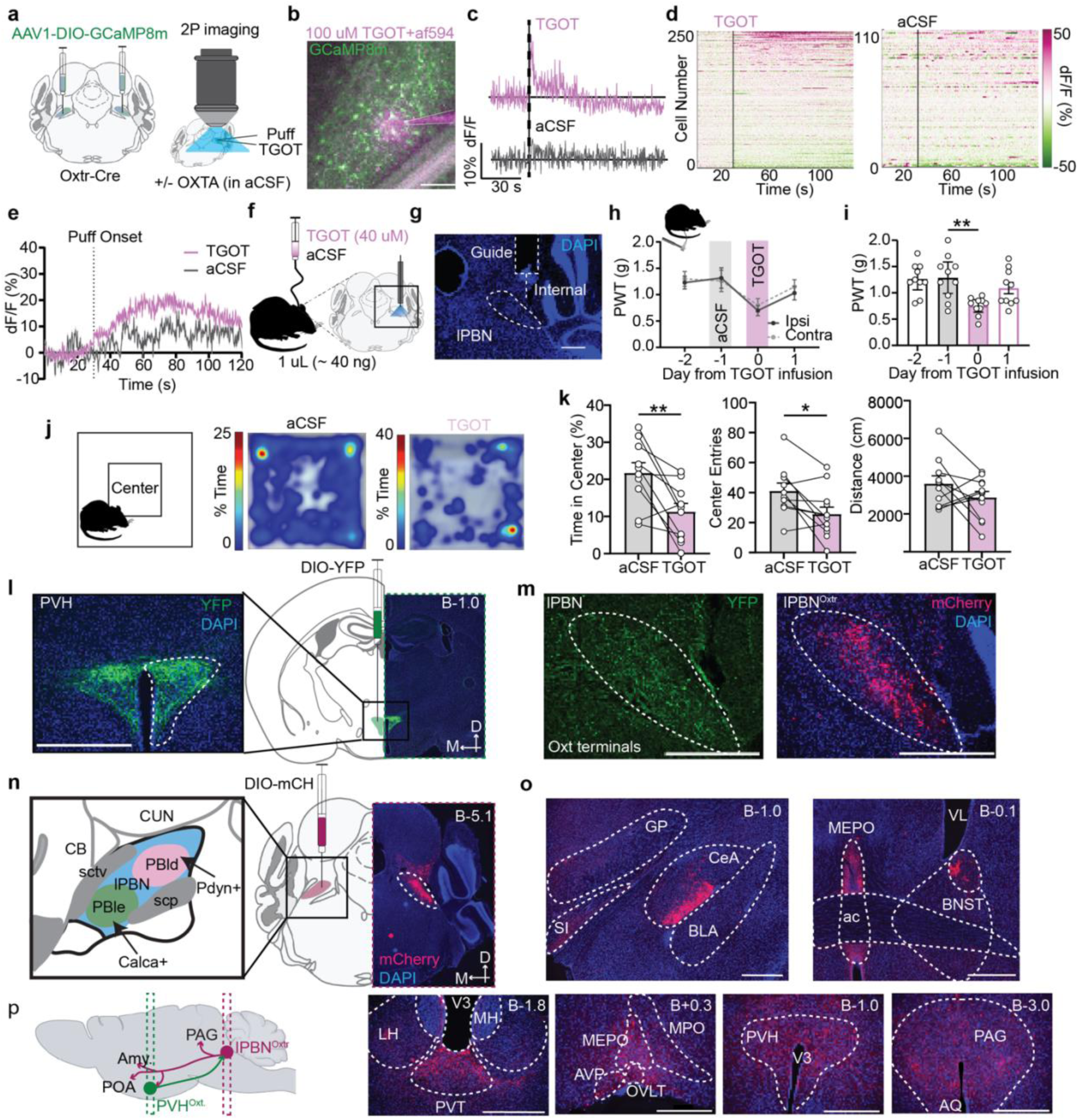
Oxytocin produces excitation of lPBN, is anxiogenic, and pronociceptive. **a.** Schematic of 2-Photon slice imaging with micropipette administration of TGOT to lPBN^Oxtr^ neurons. **b.** Example fluorescence image of GCaMP expression in lPBN and 200 ms puff of solution from the micropipette containing AlexaFluor-594 (AF594). Scale bar is 50 μm. (Left). **c.** Example cell dF/F trace (plotted as % change from baseline period (30 s)) aligned with puff from TGOT (100 μM) or aCSF vehicle control. **d.** Heat map of average cell response to 3 puff trials for TGOT (left) or aCSF (right). Line at 30 s indicates the time of puff. **e.** Response of the top 20% of cells in d to TGOT or aCSF puff. **f.** Schematic and **g.** histological example of intracranial cannula and infusion of TGOT or aCSF to the lPBN. **h.** 50% PWT (g) for the hind paw ipsilateral (solid black line) or contralateral (dotted grey line) to the implanted cannula. **i.** The average hind paw values for each mouse (one-way ANOVA with repeated measures). **j.** Open field test assay schematic with example heat maps from animals receiving either treatment. **k.** Quantification of time in center, entries to the center zone, and total distance traveled during the open field test. **l.** Schematic and histological example of viral injection carrying YFP to the PVH of Oxt-Cre mice. **m.** Example image of YFP-labeled terminals in lPBN from the mouse in l next to an example image of an Oxtr-Cre mouse injected with mCherry. **n.** Graphical depiction of anatomical subregions of lPBN with recognized gene markers (left). Histological example of viral injection carrying mCherry to the lPBN of Oxtr-Cre mice (right). **o.** Examples of mCherry+ regions of interest downstream from lPBN injection (shown in M). **p.** Schematic of anatomical findings from l-o. The anatomic location of histology images is reported as mm from Bregma at the top right of each image. All scale bars are 500 μm.

To determine the behavioral consequences of direct oxytocin receptor activation within the lPBN, we implanted guide cannulae targeting the lPBN (**Fig. 5f, g**) and administered either the selective oxytocin receptor agonist TGOT (40 µM) or aCSF as a vehicle control. Given the established role of lPBN neurons in peripheral sensitization^34,35,64^, we assessed whether TGOT infusion altered tactile sensitivity using von Frey filament application to the plantar surface. TGOT-infused animals exhibited a small but significant increase in bilateral paw sensitivity (**Fig. 5h**). This effect was largely resolved by twenty-four hours post infusion, indicating a transient but pronounced sensitizing action of lPBN^Oxtr^ activation (**Fig. 5i**). Infusion of TGOT into the lPBN was anxiogenic in the open-field test. Spatial occupancy heatmaps illustrate the thigmotactic distribution of TGOT-treated animals compared to the more exploratory patterns observed in aCSF controls (**Fig. 5j**). Relative to controls, TGOT-infused animals exhibited reduced time spent in and fewer entries to the center zone of the open field arena, a region analogous to exposed environments associated with elevated predation risk, without affecting distance traveled (**Fig. 5k**).

Having demonstrated that direct pharmacological activation of lPBN^Oxtr^ neurons is both pro-nociceptive and anxiogenic, we sought to characterize the anatomical connectivity of this population. To demonstrate that lPBN^Oxtr^ neurons receive oxytocinergic input from the hypothalamus, we induced YFP expression in the periventricular hypothalamus (PVH) of *Oxt*-Cre mice and identified YFP-positive axonal terminals within the lPBN (**Fig. 5l, m**). This is consistent with the relatively dense oxytocinergic terminal fields previously described in the brainstem and provides direct anatomical evidence that lPBN^Oxtr^ neurons are positioned to receive oxytocinergic input from the PVH^14,26^. To define the downstream projection targets of lPBN^Oxtr^ neurons, we injected *Oxtr^Cre^* mice with a virus carrying a Cre-dependent mCherry and performed whole-brain sectioning to map terminal fields (**Fig. 5n**). lPBN^Oxtr^ neurons were found to project along both the ventral and central tegmental tract output streams of the lPBN^60^, with terminal fields identified in the central nucleus of the amygdala (CeA), oval nucleus of the bed nucleus of the stria terminalis (BNST), paraventricular thalamus (PVT), median preoptic area (MEPO), periventricular hypothalamus (PVH), and the anterior periaqueductal grey (PAG) (**Fig. 5o**), consistent with recent findings^65^. Collectively, these projection targets encompass regions implicated in fluid homeostasis, pain modulation, and threat-related affect, suggesting that lPBN^Oxtr^ neurons are rooted within canonical circuits known to be relevant in a multitude of aversive and homeostatic behaviors. A schematic summarizing the input-output connectivity of lPBN^Oxtr^ neurons is provided for reference (**Fig. 5p**). The findings presented here are consistent with the interpretation that lPBN^Oxtr^ neurons are positioned to receive and respond to endogenous oxytocinergic input under physiologically relevant conditions to drive defensive behavior such as fleeing and tactile vigilance.

## DISCUSSION

The present study characterizes a relatively understudied population of *Oxtr*-expressing neurons in the lPBN and implicates oxytocin signaling at this brainstem site in the neural computations underlying both directly experienced and socially transmitted threat responses. lPBN^Oxtr^ neurons are not a functionally homogeneous extension of the broader parabrachial nociceptive system but rather a distributed population that could integrate ascending sensory signals with descending hypothalamic oxytocinergic input across a broad range of aversive contexts including acute mechanical and inflammatory pain, predator threat and the observed distress of a familiar conspecific. The behavioral consequences of manipulating this population are asymmetric: direct pharmacological activation of lPBN^Oxtrs^ is anxiogenic and pro-nociceptive, yet chemogenetic inhibition selectively disrupts the acquisition of hypersensitive states without affecting anxiety, suggesting that endogenous oxytocin signaling at this site may be engaged preferentially under conditions that demand flexible updating of defensive behavioral tone. This idea is consistent with previous findings that oxytocin signaling within the central amygdala promotes active escape behavior and inhibits freezing only when an individual animal is faced with imminent, not distant or inescapable, threats^66^. The consistent coupling of lPBN^Oxtr^ neuron activity to locomotion velocity across paradigms points to a functional organization that may reflect the interface between threat detection and the motor systems that execute appropriate avoidance responses. Collectively, these observations raise several questions that we address in turn. How does the molecular and circuit identity of lPBN^Oxtr^ neurons position them within the broader parabrachial network? How does the context of oxytocin release determine its behavioral consequences? We also consider what the sex-specific effects of lPBN^Oxtr^ neuron inhibition on affiliative proximity and pain contagion reveal about individual differences in social threat transmission, as well as how dysfunction within this system might contribute to the maladaptive hypervigilance and pain sensitization that characterize stress-related neuropsychiatric conditions.

The consistent suppression of lPBN^Oxtr^ neuron activity during behavioral arrest across multiple paradigms—foot shock, elevated plus maze, and simulated predator threat—underscores the tight coupling between this population and locomotor state. Whether inhibition during freezing reflects active suppression by upstream circuits, withdrawal of excitatory sensory drive, or an intrinsic property of the population remains to be determined. One possibility is that the activity of these neurons reflects the intensity of sensory input arriving from rapid footfalls or somatosensory surfaces during movement, and that the suppression during immobility is a consequence of reduced afferent drive rather than a behaviorally meaningful signal *per se*. Alternatively, the positive coupling between lPBN^Oxtr^ activity and velocity may reflect motivational signals related to the drive to flee anxiogenic or threatening environments, consistent with the role of the lPBN in organizing the affective and motivational components of aversive experience.

Our anatomical and molecular characterization of lPBN^Oxtr^ neurons reveals that most of this population is distinct from the well-characterized *Pdyn*- and *Calca*-expressing subpopulations of the lPBN. This is notable given prior work by which found that lPBN^Oxtr^ neurons were not responsive to tail shock when assessed using single-cell calcium imaging approaches targeted predominantly to the dorsal lateral subdivision^33^, a region with substantial overlap with *Pdyn*-expressing neurons, more strongly associated with fluid intake regulation^14,33,36,67^. The bulk photometry approach employed here, which captures the aggregate activity of the broader *Oxtr*-expressing population throughout the lPBN including the external lateral subdivision, may be sensitive to responses that are diluted or absent in targeted single-cell recordings of a more restricted dorsal subpopulation. It is also worth noting that prior work has established that modulation of liquid consumption by lPBN^Oxtr^ neurons may not be dependent on oxytocinergic projections from the hypothalamus but is instead likely driven by inputs from the nucleus of the solitary tract^14^ suggesting that functionally distinct inputs converge on molecularly overlapping but behaviorally dissociable subsets within the broader *Oxtr-*expressing population. The partial overlap of lPBN^Oxtr^ neurons with *Calca*-expressing cells is of particular interest, given the established role of CGRP-expressing lPBN neurons in peripheral sensitization and pain-related behavior^34,35^; it is possible that this overlap contributes to the nocifensive consequences of oxytocin signaling in this region.

Previous work has largely identified oxytocin as an anti-nociceptive signaling peptide, acting both through its cognate *Oxtr-*encoded receptor and TRPV1 channels, under painful conditions^68–73^. Considering this, we were somewhat surprised to see that TGOT infusion to lPBN was moderately pro-nociceptive. A subset of PVH neurons are selectively activated by pain or pain-inducing stimuli following observation of distressed social partners^59,71,74^. This supports the hypothesis that there may be a distinct output channel releasing oxytocin under pronociceptive conditions, but this has yet to be determined. Herpertz et al.,^75^ suggests that oxytocin neuron responses may be selectively enhanced by pain anticipation in humans. Our sensitivity data in CFA-observers may reflect increased vigilance toward extraneous stimuli to the paws as a form of pain-related anticipation.

A minority of lPBN^Oxtr^ neurons co-express the vesicular GABA transporter, Vgat (*Slc32a1)*, identifying them as inhibitory. The functional significance of the lPBN GABAergic subpopulation remains unclear, though their activation is antinociceptive^76^. Given the predominantly glutamatergic identity of lPBN output neurons and the inhibitory influence that local interneurons exert on parabrachial circuit dynamics, it is conceivable that Oxtr-expressing GABAergic neurons serve a circuit-level gating function, modulating the gain of excitatory lPBN output in an oxytocin-dependent manner. Whether this population is recruited under the conditions examined here, and whether it contributes to the behavioral effects of oxytocin receptor manipulation warrants dedicated future investigations.

In this study, we modeled behavioral state matching in mice wherein two animals adopt similar hypervigilant or nocifensive states despite only one—the demonstrator— directly experiencing a stressful or painful event. This phenomenon, broadly conserved across species, can prime adaptive learning^77^ and facilitate coordinated responses to environmental threats, but its underlying neural mechanisms remain incompletely resolved. The neuro-physiological consequences of aversive internal states are broadly similar in demonstrators and observers, and the involvement of frontal-cortical and amygdala circuits in such observational transfer has been demonstrated across humans, non-human primates, and rodents^38,43,46,78–81^. Notably, frontal-cortical activity appears necessary for the acquisition of aversive observational behavior but is dispensable for its maintenance, suggesting that once established, observer hypersensitivity may be sustained by peripheral sensitization at the level of the spinal cord^82^. The present findings identify lPBN^Oxtr^ neurons as a brainstem node that similarly contributes to the acquisition, but not the maintenance, of observer-induced mechanical hypersensitivity, extending the circuitry implicated in socially transmitted affective states to a region with privileged access to ascending nociceptive and interoceptive signals.

Human studies demonstrate that high empathy for a partner in pain enhances one’s own pain sensitivity and vulnerability to negative affect^83–85^, while over-attribution of negative valence to neutral social cues is thought to underlie several social anxiety disorders^53^. Conversely, impaired processing of and responsiveness to social cues, a core feature of autism spectrum disorder, carries its own consequences for well-being and mental health^54^. Social learning and the transmission of affective states are thus bidirectionally relevant to neuropsychiatric disease and understanding how the brain represents the internal state of a social partner is a key step toward understanding how such processes become dysregulated. The high behavioral variance observed in both human and rodent studies of social transmission may be partially explained by the epigenetic tuning of oxytocin receptor distribution, which is individual-specific and sensitive to developmental, environmental and experiential factors^1,3^. This variability is reflected in our own data, where observer responses to both CFA and foot shock paradigms were more heterogeneous than those of demonstrators, and where sex-specific differences in affiliative proximity during chemogenetic inhibition experiments were observed without corresponding differences in the magnitude of transmitted hypersensitivity.

### Limitations

Our experiments demonstrate that direct infusion of the selective Oxtr agonist TGOT into the lPBN is anxiogenic in the open field, yet our neural manipulations did not alter anxiety in the elevated plus maze. This apparent discrepancy likely reflects the limitations inherent to pharmacological activation studies, in which the volume of infusate, the concentration of agonist, and the spatial spread of drug delivery impose constraints on interpretation. A single infusion of 1 µL places a substantial quantity of TGOT within and potentially beyond the lPBN, raising the possibility of off-target receptor engagement in adjacent structures. Alternatively, it is possible that TGOT is binding Vasopressin receptors at the higher concentrations used. Though TGOT is 1000-fold more selective for oxytocin than vasopressin receptors, vasopressin receptor activation is anxiogenic in other brain regions^3,86^. We attempted small, 250 nL, volume infusions which were confounded by technical limitations of our infusion technique but showed a trend toward inducing tactile sensitivity without impacting anxiety metrics in the elevated plus maze (**Extended Fig. 5**). Future studies employing more spatially restricted pharmacological approaches or real-time measures of oxytocin release will be necessary to resolve the relationship between endogenous oxytocinergic signaling, locomotion, and anxiety-related behavior at this site. It will also be of interest to examine threat avoidance behavior more directly, given the consistent positive coupling between lPBN^Oxtr^ neuron activity and velocity and the established role of the lPBN in escape and avoidance^31,32,64,87–91^.

A technically important consideration in interpreting the fiber photometry data from social observation experiments is the difficulty of detecting stimulus-evoked changes in lPBN^Oxtr^ activity against a shifting fluorescence baseline across sessions. Because baseline GCaMP fluorescence can differ substantially between a pre-CFA and post-CFA session, likely reflecting an elevated tonic activity state of the population in the presence of ongoing inflammatory input, standard normalization approaches may underestimate the true magnitude of phasic responses to von Frey filament application or other discrete stimuli in the pain state. This technical limitation could account for our inability to detect a significant difference in the neural response to filament application as a function of pain induction or observation, despite the robust behavioral differences observed across these conditions. Development of normalization strategies that account for session-level shifts in baseline fluorescence will be an important methodological advance for future work in this area.

## ACKNOWLEDGEMENTS

We thank Lucy Anastas, Susan Phelps, and Kit Mandeville for mouse colony maintenance, Hadiya Amjad, Kishan Sood, Rachel Felix, and Michael Boakye-Ansah for assistance with behavioral experiments and manual scoring. We would like to thank Drs. Azra Suko, Selena Schrieber, and Larry Zweifel for their help with virus production. We also thank our colleagues for useful discussions.

## AUTHOR CONTRIBUTIONS

**EJ:** Conceptualization (lead); Investigation (lead); Writing – original draft (lead). **JP**: Histological assistance and quantification. **SK**: Behavioral experiments and quantification, **MRB** and **RDP**: research support, advice and editing.

## METHODS

All experiments were approved by the University of Washington Institutional Animal Care and Use Committee and were performed in accordance with the guidelines described in the US National Institutes of Health Guide for the Care and Use of Laboratory Animals. *Oxtr^Cre^* and *Oxt^Cre^* mice were obtained from Jax (strain # 031303, 024234, respectively) and bred in house to maintain heterozygous Cre-expressing lines for experimental studies. All mice were backcrossed onto a C57BL/6 background. Both male and female mice were used for all experiments, aged 8-12 weeks at the start of experimental procedures and no more than 20 weeks at the end of experimental procedures. Mice were housed on a 12-h light/dark cycle (lights on 05:00-17:00) at ∼22°C with food and water available *ad libitum*.

### Stereotaxic surgery

Mice were anesthetized initially with 4% isoflurane and then maintained at 1-2% at a flow rate of 1 L/min. After anesthesia induction mice were placed on a stereotaxic frame (David Kopf Instruments). All stereotaxic coordinates are in mm relative to the location of Bregma following medial/lateral and dorsal ventral leveling. Viral injections were performed with the Nanoject II system unilaterally for imaging and bilaterally for chemogenetic or mutagenesis experiments into the PBN (anterior-posterior: −5.1, medial-lateral: +/−1.6, dorsal-ventral: +/-3.2-3.3) at a rate of 0.1 μL/min for 2.5 min for a total of 0.25 μL. All injections to the PVH were performed bilaterally (anterior-posterior: −0.5, medial-lateral: +/−0.3, dorsal-ventral: +/-4.6). Following viral injection, injection needles were maintained in place for 4 minutes before removal at each injection site. For optic fiber or intracranial cannulae experiments, cables were implanted immediately dorsal (anterior-posterior: −5.1, medial-lateral: +/−1.6, dorsal-ventral: +/-2.8 (guide cannulae) 3.2 (optic fiber: 4 00 µm core, 1.25 mm ferrule, Doric) to the viral injection site. To secure the implants to the skull, C&B Metabond (Parkell) and dental cement were used. At the end of each experiment animals were euthanized with phenobarbital and the brain was extracted for histological evaluation of viral placement and fiber optic placement. Animals were allowed to recover for 4 weeks following viral injection and 1 week following intracranial cannula implantations to limit clogging of the guide cannula with fibrotic tissue.

### Behavioral Assays

#### Paw Withdrawal Measurements

The von Frey, tactile-sensitivity assay was performed using the ascending application method. Each animal was placed in a 11.5-cm by 7.5-cm chamber with a wire mesh floor. Animals were acclimated to the chamber for 30 min before the assay began. Filaments (BioSeb) were applied to a 45-degree angle at the plantar surface of each hind paw for 1-2 seconds a total of 5 times. Filament application began with a 0.16-g filament and ended after two consecutive filaments elicit a paw-withdrawal on 3 or more out of 5 applications. A positive paw-withdrawal response was accompanied by licking, flicking, or immediate guarding of the stimulated paw. For photometry experiments, animals were habituated to the patch cord and von Frey testing chambers before recording began. During photometry recordings, animals were connected to the patch cord and allowed to habituate to the von Frey chambers for 10 minutes. Each recording consisted of a 3 minute baseline period followed by 8 filament applications at 0.6 g then 2.0 g with an ITI of roughly 100 s. Each trial alternated the targeted hind paw such that each paw received 4 applications at each weight. Pin application trials consisted of 2 trials per hind paw and were always completed at the end of the recording session to reduce sensitization. Pin trials did not draw blood or induce persistent inflammation.

#### Open Field

Animals were placed in the center of an illuminated square open field chamber (40 cm by 40 cm) and were then allowed to freely explore for 10 minutes before being removed the from the chamber and placed in back in their home cages. The center area was determined as a square in the center of the field that was 50% smaller than the surrounding area. The center point of the animal was tracked using Ethovision Software. The center point of the animal had to cross 2 cm beyond the zone edge to be considered within the center zone.

#### Predator Chase (RoboBug)

Animals were attached by patch cord to the Fiber Photometry Cart (Tdt RZX) and allowed to acclimate in their home cage for 5 minutes. They were then placed in the center of an illuminated square open field chamber (50 cm by 50 cm) and were allowed to freely explore for 10 minutes with a stationary robotic toy bug (Hexbug Red Spider Micro Robotic Creature Hex Bug) in one corner. Following the 10-min baseline period, the robotic bug moved directly toward the mouse for 10 s before retreating to a corner and remaining in place for a minimum of 2 min. Each trial was experimenter-initiated when the bug and mouse were roughly caddy-corner to one another. Each mouse experienced 5 trials of predator-like approach. Videos of each session were captured with USB camera and a personal computer. The center point of the animal was tracked using Ethovision Software (XT18). Velocity measures were calculated using a moving average over 10 frames throughout the session.

#### Elevated Plus Maze (EPM)

The custom-made EPM consisted of two sets of crossed arms (two arms enclosed by 30 cm tall transparent plexiglass, two arms open), each 40 cm long and 8 cm wide, set 70 cm above floor. Animals were placed in the center of the EPM facing alternating closed arms. Animals were then allowed to freely explore for 10 minutes before being removed the from the apparatus and placed in back in their home cages. Videos of each session were captured with USB camera and a personal computer. The center point of the animal was tracked using Ethovision Software (XT18) and the two open and two closed zones were combined, respectively, for final calculations. The center point of the animal had to cross 2 cm beyond the zone edge to be considered within each zone. Velocity measures were calculated using a moving average over 10 frames throughout the session.

#### Foot Shock and Auditory Fear conditioning

The fear-conditioning chamber was a square arena (25 × 25 cm) with metal walls, two speakers attached on opposite walls, and a metal grid floor that consisted of a circuit board that delivers electrical shock (Coulbourn Instruments). A USB camera was connected to the personal computer and recorded the data. Animals were habituated to the chambers a minimum of 3 days for 20 minutes and allowed a 5-minute baseline period within the chambers where no shock or auditory stimulus was delivered in all subsequent testing days. White noise was played in each box to minimize auditory contamination from neighboring boxes. Foot shocks were always delivered at 0.5 mA for 1 s.

#### Observational fear conditioning

Shock chambers were split in two with custom made L shaped clear plexiglass dividers with perforations of 1 cm diameter along the wall side. Animals were habituated in pairs to the split chamber for a minimum of 3 20-minute sessions on consecutive days. Animals were then subjected to a conditioned stimulus (CS) alone day (4 CS trials; Day 1), two conditioning sessions consisting of 10 trials where the CS co-terminated with a 1 s foot shock (0.5 mA; Days 2 and 3), and a final CS only day (4 CS trials) where one animal was in the chamber but the plexiglass floor remained (Day 4). The demonstrator (animal who received shocks during Days 2 and 3) was placed on the plexiglass floor for Day 4 and the observer (who observed foot shocks on Days 2 and 3) was placed on the grid floor for Day 4. Essentially this swapped the context for each individual in the conditioned pair. The CS consisted of a 3 kHz, 10 s, 80 dB constant tone played at random intervals, with an average inter-trial interval (ITI) of 90 s.

All trials were recorded by a USB camera attached to the personal computer and the time spent freezing, defined as immobility (patch cords excluded in photometry experiments), was analyzed post-hoc using EzTrack freezing software which relies on pixel change above the threshold of an empty chamber (grayscale difference value of 10-13 depending on box conditions). The threshold for freezing was determined by eye as a maximum motion value of 60 (a.u.) for a minimum of 30 frames or 1 second. To validate ezTrack’s fidelity in calculating freezing epochs, two individual researchers blind to the experimental conditions scored 3-90 second periods for freezing behavior (**Extended data, Fig 1**). The percent of time spent freezing was then compared between researchers (average difference of ∼2%) and between ezTrack-calculated values (average difference of 1.4 and -0.8%).

#### Observation of Inflammatory Pain

These experiments with cross housing were completed as described by^38,39^. For animals that remained pair housed for the duration of the experiments, experiments went as follows. Animals were handled and habituated to the von Frey apparatus and i.p. injections for 1 week prior to the start of behavioral experiments. Female mice were pair housed with female siblings if available or age-matched female wildtype mice for a minimum of 4 weeks after surgery and before behavioral testing. Male animals were always pair housed with a sibling. Male demonstrator mice did not receive viral injections but did receive surgical incisions and sutures to minimize fighting and disruption of surgical sites We video recorded the first ten and last ten minutes of this interaction period. The videos were then processed using Ethovision and the two-animal tracks were corrected by hand for identity swapping or fidelity issues. We then used the distance between the center points to assess proximity between the two mice throughout the two ten-minute periods. CFA injections (20uL) were done in the subplantar area by inserting a 31-gauge needle between the toes and into the ventral paw pad. Controls were restrained in same way and poked with a pen but received no subplantar injection to minimize any lasting ‘pain’ component or scent of blood.

### Pharmacological injections

Clozapine N-oxide (Sigma Aldrich), nitroglycerin (American Reagent, Inc), OXTA (L368,899, Med Chem Express) and saline or vehicle controls were injected at 10 mL/kg body weight. CNO was administered at 3 mg/kg for hM4Di experiments, nitroglycerin at 10 mg/kg, L368,899 hydrochloride at 10 mg/kg), and saline at 0.9% sodium chloride; vehicle for nitroglycerin was 6% propylene glycol 6% ethanol and saline. TGOT (Med Chem Express) was prepared at 4 mM in purified, sterile water and diluted in aCSF to working concentrations. aCSF vehicle controls for TGOT contain the same amount of purified, sterile water as TGOT preparations.

### Immunohistochemistry

Mice were deeply anesthetized with Euthasol (0.25 mL, i.p.) and perfused transcardially with phosphate-buffered saline (PBS) followed by 4% paraformaldehyde (PFA, Electron Microscopy Sciences) in PBS. Brains were post-fixed overnight in 4% PFA at 4°C, cryoprotected in 30% sucrose, frozen in OCT compound and stored at −80°C. Coronal sections (40 μm) were cut on a cryostat (Leica Microsystems) and collected in cold PBS. For immunohistochemistry experiments, sections were washed three times in PBS with 0.2% Triton X-100 (PBST) for 5 min and incubated in blocking solution (3% normal donkey serum in PBST) for 90 minutes at room temperature. Sections were incubated overnight at 4°C in PBS with primary antibodies including: chicken-*anti*-GFP (1:2000, Abcam, ab 13970) or rabbit-*anti*-dsRed (1:1000, Takara, ab 632496). After 3 washes in PBS, sections were incubated for 1 h in PBS with secondary antibodies: Alexa Fluor 488 donkey anti-chicken or Alexa Fluor 594 donkey anti-rabbit (1:500, Jackson ImmunoResearch). Tissue was washed 3 times in PBS, mounted onto glass slides, and cover slipped with Fluoromount-G containing DAPI (Southern Biotech). Fluorescent images were acquired using a Keyence BZ-X700 microscope. Images were minimally processed using ImageJ software (NIH) to enhance brightness and contrast for optimal representation of the data. All digital images were processed in the same way between experimental conditions to avoid artificial manipulation between different datasets.

### RNAscope *in situ* hybridization

Mice were anesthetized with Euthasol (0.25 mL, i.p.) then decapitated. Brains were rapidly frozen on crushed Dry Ice. Coronal sections (20 μm) were cut on a cryostat (Leica Microsystems), mounted onto glass slides, and stored at −80°C. RNAscope fluorescent multiplex assay was performed following the manufacturer’s protocols. Several levels of the PBN were imaged for each animal using a Keyence BZ-X710 microscope. Using the superior cerebellar peduncle (scp) as the center point, images were acquired at 20× in a 3 × 3 grid then stacked and stitched together using Fiji. Background fluorescence was subtracted using the image calculator function in FIJI. Images of probe staining within the four-channel sets minimally processed to enhance brightness and contrast for optimal representation of the data. PBN anatomy was estimated using fiber-tract location and general structure, relying on the presence of the scp and the ventral spinocerebellar tract (sctv) as well cerebellar morphology to determine the regions of interest as lPBN. Images were imported into QuPath^92^ and a region of interest was drawn over the lateral PBN using the surrounding fiber tracts and brain structure as a guide. RNA expression was quantified by thresholding using the subcellular detection function in QuPath.

### 2-photon slice imaging

Mice were deeply anesthetized with Euthasol (0.25 mL, i.p.) and intracardially perfused with ice-cold cutting solution containing (in mM): 92 N-methyl-D-glucamine, 25 D-glucose, 2.5 KCl, 10 MgSO_4_, 1.25 NaH_2_PO_4_, 30 NaHCO_3_, 0.5 CaCl_2_, 20 HEPES, 2 thiourea, 5 Na-ascorbate, 3 Na-pyruvate. Brains were quickly removed after perfusion and 250-μm coronal slices were prepared (Leica VT1200) in the same ice-cold solution. Brain slices were kept in the cutting solution at 33°C for 10 min and then transferred to a room temperature recovery solution containing (in mM): 13 D-glucose, 124 NaCl, 2.5 KCl, 2 MgSO_4_, 1.25 NaH_2_PO_4_, 24 NaHCO_3_, 2 CaCl_2_, 5 HEPES for at least 1 h. Slices were individually transferred to 33°C artificial cerebral spinal fluid containing (in mM): 11 D-glucose, 126 NaCl, 2.5 KCl, 1.2 NaH_2_PO_4_, 26 NaHCO_3_, 2.4 CaCl_2_, 1.2 MgCl_2_ for recording. All solutions were saturated with 95% O2/5% CO2 and adjusted to pH 7.3–7.4. TGOT or aCSF vehicle control solutions were applied using a pressure system where carboxide gas delivered roughly 0.07 mmHg of pressure for 200 ms through plastic tubing connected to a micropipette (ID X, OD 1.2 mm) resulting in the expulsion of approximately 0.2 uL. The micropipette was positioned near GCaMP expressing neurons using a micromanipulator. Each recorded slice was bathed in aCSF containing AMPA (0.2 mM) as a positive control. Only AMPA-responsive (a mean fluorescence after bath application of 1.65 times the standard deviation of the baseline fluorescence) neurons were further analyzed. Neuropil-subtracted fluorescence traces were gathered for each puff trial and normalized to the 30 s baseline period before puff delivery. For each neuron, we averaged the trace of 3 consecutive puff trials. The neurons were then sorted by their mean dF/F response in the 60 s following puff delivery. The top 20% of neurons in each group were then averaged and compared across TGOT and vehicle puff conditions.

### Cannula infusions

Mice were given 1 week to recover from cannula implantation before being habituated to handling. The animals were handled every day for 1 week before first infusion. For intracranial infusion, TGOT (40 mM; 10 ng/uL) or aCSF were loaded into an injection cannula (ProTech) connected to a microinjector (1 μl total volume). TGOT was dissolved in sterile water to 0.2 mM then diluted in aCSF (same as slice imaging) to 40 mM. aCSF vehicle controls contained the same volume of sterile water as TGOT preparations. A total of 1 μL of solution was infused at a rate of 8 nL/s (over 2 min). Animals were kept connected to the cannula for an additional 3-5 min to allow for stable diffusion of the solution away from the tip. Afterward, the internal cannula was removed and replaced with a dummy cannula to protect the guide cannula opening and maintain adequate patency. Animals were returned to their home cage for a minimum of 15 minutes before additional behavioral testing.

### Fiber photometry recordings and analysis

Fiber photometry studies were completed as described previously ^93^. In brief, GCaMP6m fluorescence was excited using a 470-nm LED (Ca^2+^-dependent signal) and a 405-nm LED (isosbestic control, Ca^2+^-independent signal). LED intensities were set to 30 µW at the optic fiber tip. GCaMP6m emissions were filtered (525 ± 25 nm), detected with a photoreceiver, and recorded by a real-time processor (Tucker Davis Technologies). Fiber photometry data were analyzed as described^93,94^. In brief, custom MATLAB scripts were used to normalize signal by detrending decay from bleaching then dividing by a linear least-square fit of the isosbestic trace scaled to the signal. The processed traces were extracted in windows surrounding the onset of relevant behavioral events (tail lift, shock, zone entry, velocity change, or post-hoc manually scored behavior), z-scored relative to the mean and standard deviation of the full recording session and normalized to the pre-event baseline by subtraction. The trials were then averaged within subject, then across subjects. Grand mean and standard error of the mean are plotted across subjects.

### Statistical analyses

Statistical analyses were performed as indicated in GraphPad Prism 9 and MATLAB (MathWorks). All data are expressed as mean ± SEM unless otherwise specified. * p <0.05, ** p<0.01, *** p <.001, **** p<0.0001.

## EXTENDED DATA FIGURES

**Extended Figure 1.**
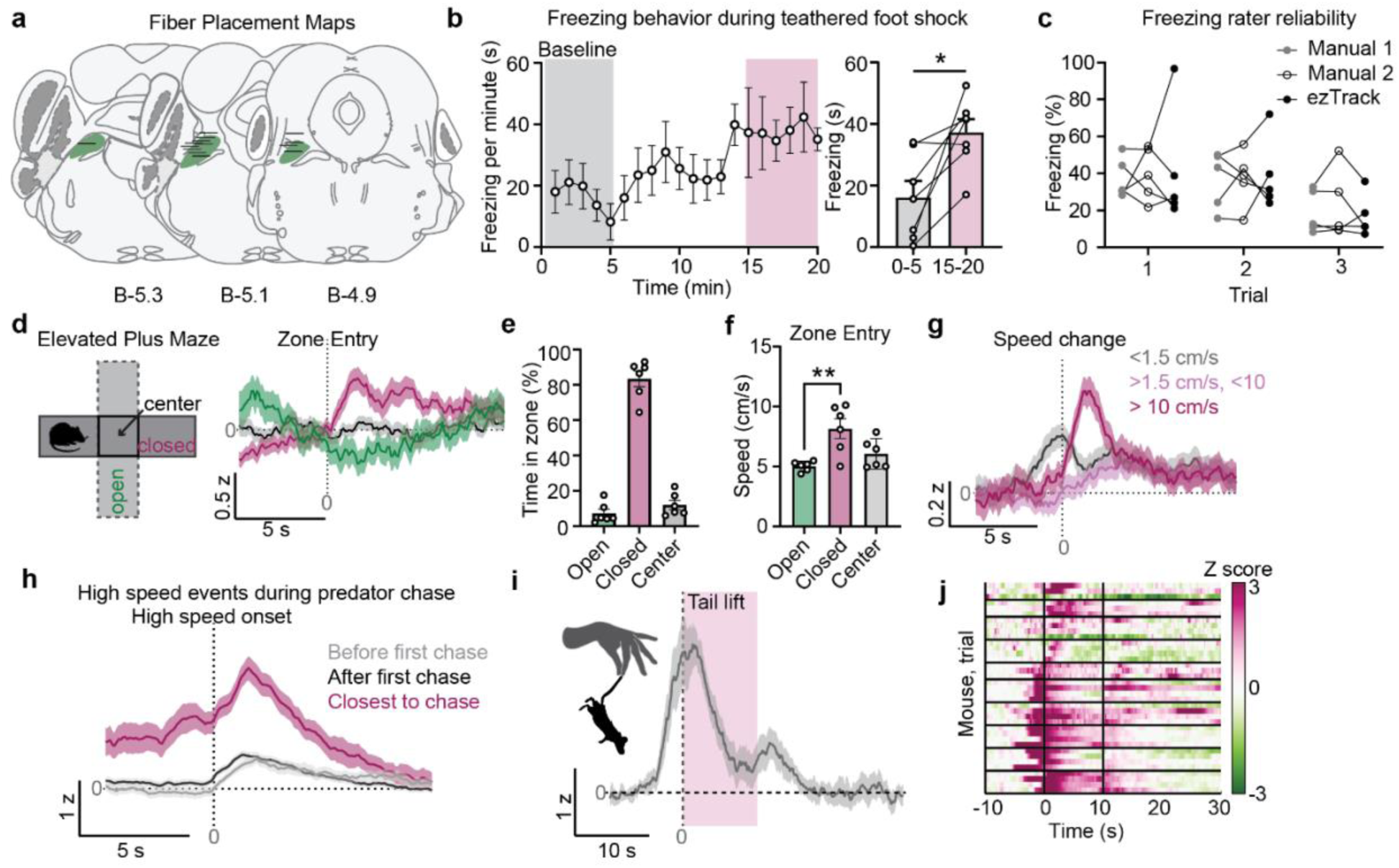
Fiber placements are restricted to lPBN and lPBN^Oxtr^ neuron activity scales with locomotion speed across anxiogenic contexts. **a.** Schematic showing the anatomical location of the approximate middle of the fiber tip for animals included in photometry analysis. **b.** Freezing values per minute averaged across seven mice with 10 shock trials (left). Quantification of freezing in the first 5 minutes before foot shock (baseline) and last 5 minutes of the end of the session (right). 10 foot shocks were delivered with a variable ITI between 30 and 120 s with an average of 90 s. **c.** Comparison of manually scored freezing with automatically detected freezing. Two independent and trained experimenters manually scored the same 30 s trials around foot shock for freezing in 4 mice (3 each). The range of detected freezing between each scorer was not significantly different in our experimental set up. **d.** Peri-event averages of calcium fluorescence aligned to entry of the closed, open, or center zones in the EPM (n =6). **e.** Quantification of the time in each zone in the EPM over a 10-minute session. **f.** Quantification of the speed at the time of entry to each zone in the EPM. **g.** Peri-event averages of calcium fluorescence aligned to speed change onset and binned by speed as detected by Ethovision. **h.** Peri-event averages of calcium fluorescence aligned to onset of high-speed events during predator chase session. Neural responses averaged over high-speed events during the baseline period (10 min, gray), after the first chase (black), or immediately following the chase (magenta). **i.** Peri-event averages of calcium fluorescence aligned to experimenter tail lift and 10 s suspension (n = 10). **j.** Heat map depicting each trial separated by individual animal. Each row depicts a trial with the earliest trial toward the bottom.

**Extended Figure 2.**
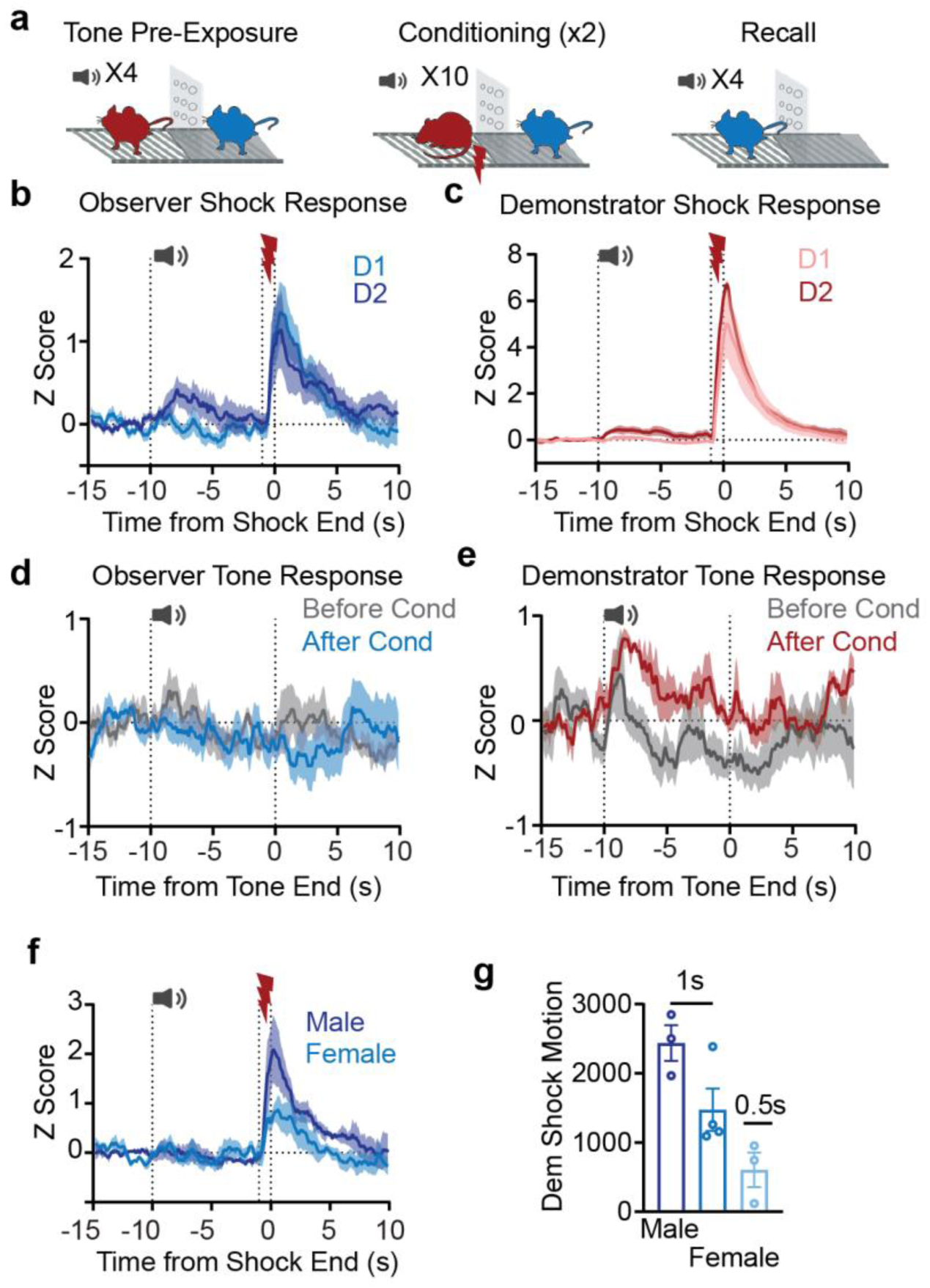
Observer lPBN^Oxtr^ neuron responses to the CS+ strengthen across conditioning sessions but do not persist to CS+ recall in the absence of the demonstrator. **a.** Schematic representation of observational fear conditioning sessions. After habituation, animals experienced 4 CS+ trials without foot shock (before conditioning). Each pair underwent 2 conditioning days where 10 CS+ trials co-terminated with a 1 s foot shock. The final recording session consisted of each animal alone in the chamber with 4 CS+ trials (after conditioning). **b.** Peri-event averages of observer calcium fluorescence aligned to the end of demonstrator foot shock in each conditioning session (D1 vs D2). **c.** Same as b but depicts demonstrator calcium fluorescence. **d.** Peri-event averages of observer calcium fluorescence aligned to the end of the CS+ tone before and after conditioning. **e.** Same as d but depicts demonstrator calcium fluorescence to the CS+ tone before and after conditioning. **f.** Peri-event averages of calcium fluorescence aligned to shock end in the 1 s shock observational fear conditioning experiment with observers separated by sex**. g.** Demonstrator shock motion separated by sex and shock length with males showing the highest shock response and females experiencing 0.5 s showing the lowest shock response.

**Extended Figure 3.**
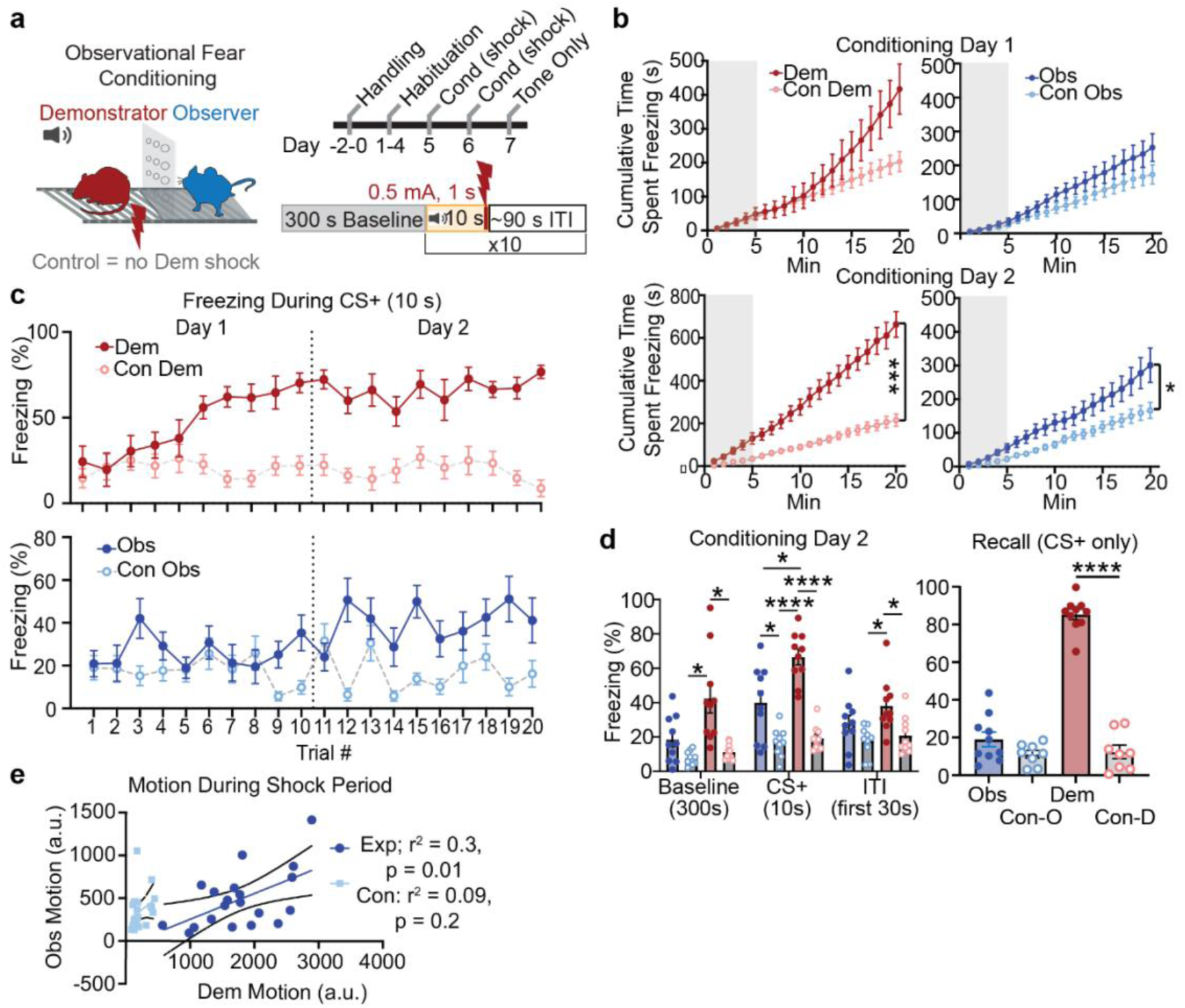
Conditioning strengthens observer freezing to witnessed shock but conditioned responses to the CS+ alone are not sustained at recall. **a.** schematic of observational fear conditioning experiments in behavioral animals (no photometry or tethering). Control animals underwent the same procedures without foot shock delivery on conditioning sessions. **b.** Cumulative time spent freezing over the conditioning sessions (top day 1, bottom day 2). Demonstrators and controls (shock vs no shock) shown in red on the left. Observers and controls (observed shock vs observed no shock) shown in blue on the right. **c.** Freezing responses during the 10 s CS+ tone throughout the conditioning sessions (trials 1-10 on day 1, 11-20 on day 2). **d.** Average freezing values to the baseline, CS+, and first 30 s of the ITI period for each group on conditioning day 2 (left). Average freezing values to the CS+ during the final tone only, recall session for each group (right). **e.** Observer motion during the final second of the CS+ on conditioning days plotted as a function of the demonstrator motion during the same epoch. Experimental pairs where shock was delivered during this time showed a significant correlation between the two animals’ motion. Control pairs’ motion was not correlated.

**Extended Figure 4.**
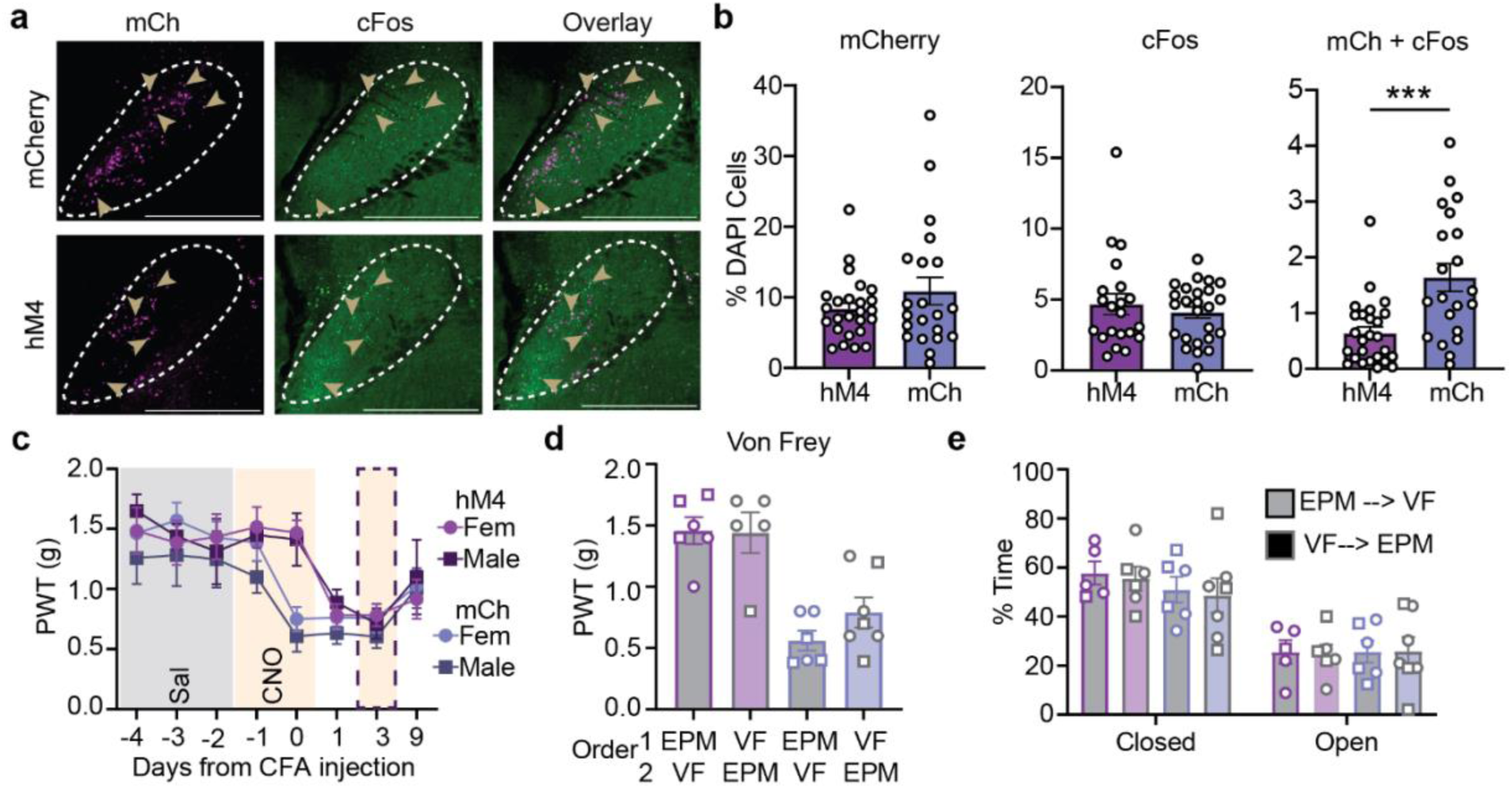
Establishing hM4Di inhibition and analysis of behavioral experiments shown in. Fig. 4**. a**. immunohistochemistry of cFos in lPBN 90 minutes after CNO injection (i.p., 3 mg/ kg) in mCherry controls or hM4Di (hM4) injected animals. **b.** quantification of mCherry, cFos, and co-expressing neurons in lPBN. **c.** Paw withdrawal thresholds (PWT) in mCherry or hM4Di CFA-observer animals split by sex. **d.** PWT in CFA-observer animals split by the order of behavioral experiments following social interaction. Grey bars with colored borders denote animals that underwent EPM before von Frey testing. Colored bars with gray borders denote animals that underwent von Frey testing before EPM testing. **e.** same as d but for EPM results-namely the percent of time spent in the open or closed arms of the EPM apparatus.

**Extended Figure 5.**
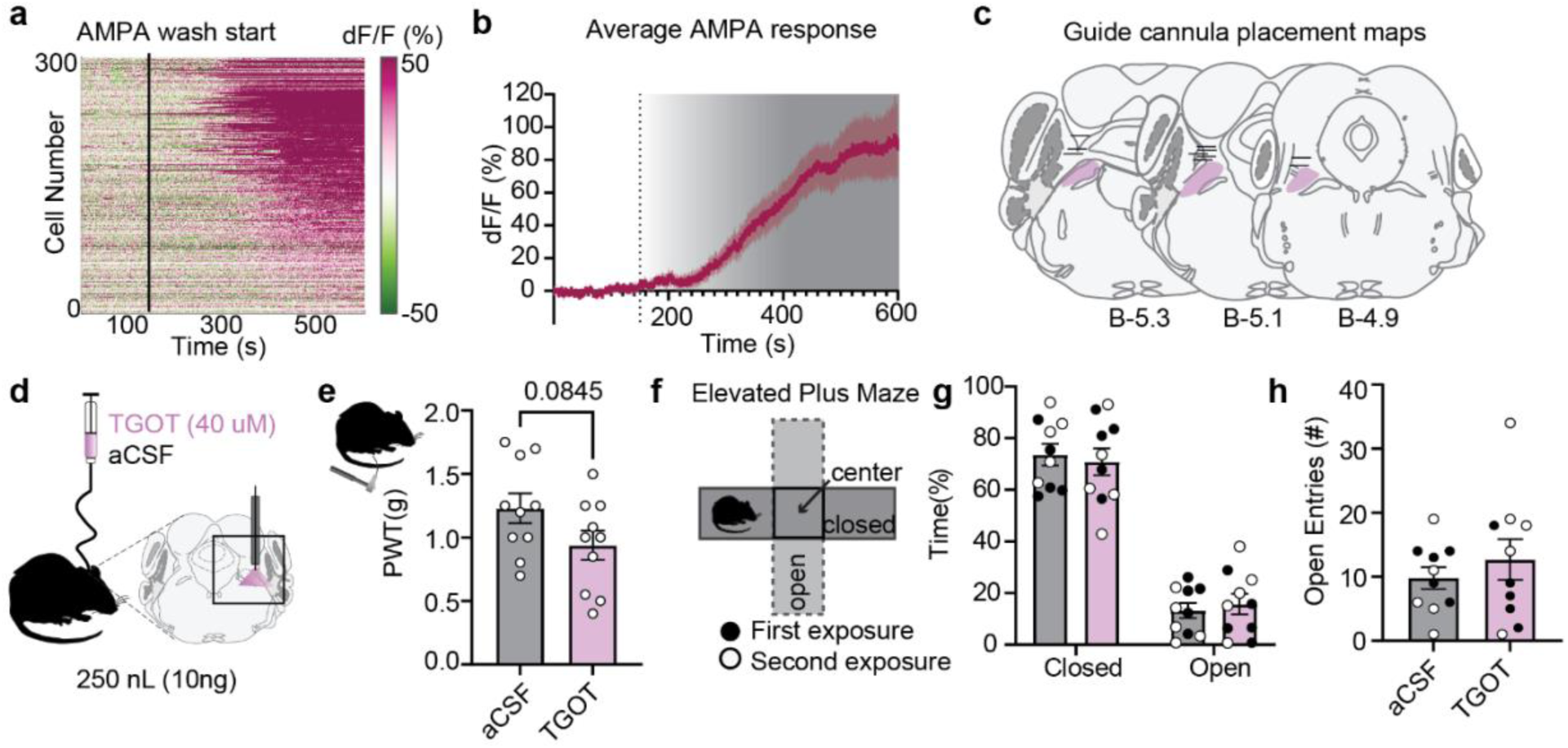
lPBN^Oxtr^ neurons are activated by bath application of AMPA and small volume cannula infusions show limited effects. **a.** Heatmap depicting lPBN^Oxtr^ neurons recorded in 2P slice experiments during the bath application of AMPA as a positive control at the end of recording sessions. Line at 150 s depicts the onset of AMPA+ aCSF bath application. **b.** Average dF/F (percent of baseline fluorescence) response of the cells shown in a. **c.** Schematic showing the anatomical location of the approximate middle of the guide cannula tip for animals included in Fig. 5. **d.** Schematic showing small volume (250 nL) cannula infusion. **e.** PWT of animals after small volume infusion of aCSF or TGOT. **f.** Cannulated animals underwent EPM testing following small volume infusion of aCSF and TGOT infusion. Treatments were counterbalanced such that each animal experienced the EPM twice, once with each treatment. Subsequent data in g and h are shown so that animals who received aCSF or TGOT on first exposure are filled circles and second exposure are open circles. **g.** Percent time spent in the open or closed arms of the EPM after infusion of aCSF (gray) or TGOT (pink). **h.** The number of open arm entries in the EPM after small volume infusion.

## REFERENCES

1. Froemke, R. C. & Young, L. J. Oxytocin, Neural Plasticity, and Social Behavior. Annual Review of Neuroscience 44, 359–381 (2021).

2. Grieb, Z. A. et al. Sex-dependent regulation of social avoidance by oxytocin signaling in the ventral tegmental area. Behav Brain Res 462, 114881 (2024).

3. Carter, C. S. The Oxytocin-Vasopressin Pathway in the Context of Love and Fear. Front Endocrinol (Lausanne) 8, 356 (2017).

4. Insel, T. R. & Shapiro, L. E. Oxytocin receptor distribution reflects social organization in monogamous and polygamous voles. Proc Natl Acad Sci U S A 89, 5981–5985 (1992).

5. Zelmanoff, D. D. et al. Oxytocin signaling regulates maternally directed behavior during early life. Science 389, eado5609 (2025).

6. He, Z. et al. Paraventricular Nucleus Oxytocin Subsystems Promote Active Paternal Behaviors in Mandarin Voles. J. Neurosci. 41, 6699–6713 (2021).

7. Menon, R. & Neumann, I. D. Detection, processing and reinforcement of social cues: regulation by the oxytocin system. Nat. Rev. Neurosci. 24, 761–777 (2023).

8. Triana-Del Rio, R., et al. The modulation of emotional and social behaviors by oxytocin signaling in limbic network. Frontiers in Molecular Neuroscience 15, (2022).

9. Eliava, M. et al. A New Population of Parvocellular Oxytocin Neurons Controlling Magnocellular Neuron Activity and Inflammatory Pain Processing. Neuron 89, 1291–1304 (2016).

10. Jiang, W.-Q., Bao, L.-L., Sun, F.-J., Liu, X.-L. & Yang, J. Oxytocin in the periaqueductal gray mainly comes form the hypothalamic supraoptic nucleus to participate in pain modulation. Peptides 121, 170153 (2019).

11. Neugebauer, V. et al. Amygdala, neuropeptides, and chronic pain-related affective behaviors. Neuropharmacology 170, 108052 (2020).

12. Liu, S. et al. Divergent brainstem opioidergic pathways that coordinate breathing with pain and emotions. Neuron 0, (2021).

13. Li, J. & Ryabinin, A. E. Oxytocin Receptors in the Mouse Centrally-projecting Edinger-Westphal Nucleus and their Potential Functional Significance for Thermoregulation. Neuroscience 498, 93–104 (2022).

14. Ryan, P. J., Ross, S. I., Campos, C. A., Derkach, V. A. & Palmiter, R. D. Oxytocin-receptor-expressing neurons in the parabrachial nucleus regulate fluid intake. Nat Neurosci 20, 1722–1733 (2017).

15. Rea, J. J. et al. Oxytocin neurons in the paraventricular and supraoptic hypothalamic nuclei bidirectionally modulate food intake. Mol Metab 100, 102220 (2025).

16. Wald, H. S. et al. NTS and VTA oxytocin reduces food motivation and food seeking. Am J Physiol Regul Integr Comp Physiol 319, R673–R683 (2020).

17. Vandendoren, M. et al. Oxytocin neurons signal state-dependent transitions to thermogenesis and behavioral arousal in social and non-social settings. 2024.10.22.619715 Preprint at 10.1101/2024.10.22.619715 (2024).

18. Amico, J. A., Miedlar, J. A., Cai, H.-M. & Vollmer, R. R. Oxytocin knockout mice: a model for studying stress-related and ingestive behaviours. Prog Brain Res 170, 53– 64 (2008).

19. Araujo, E. V. et al. Oxytocinergic signalling in the respiratory parafacial region increases the activity of chemosensitive neurons and respiratory output. J Physiol 603, 5827–5849 (2025).

20. Ferretti, V. et al. Oxytocin Signaling in the Central Amygdala Modulates Emotion Discrimination in Mice. Current Biology 29, 1938–1953.e6 (2019).

21. Pisansky, M. T., Hanson, L. R., Gottesman, I. I. & Gewirtz, J. C. Oxytocin enhances observational fear in mice. Nat Commun 8, 2102 (2017).

22. Carcea, I. et al. Oxytocin neurons enable social transmission of maternal behaviour. Nature 596, 553–557 (2021).

23. Pilgeram, N. R. et al. Oxytocin receptor antagonism during early vocal learning reduces song preference and imitation in zebra finches. Sci Rep 13, 6627 (2023).

24. Osakada, T. et al. A dedicated hypothalamic oxytocin circuit controls aversive social learning. Nature https://doi.org/10.1038/s41586-023-06958-w (2024) doi:10.1038/s41586-023-06958-w.

25. Akinrinade, I. et al. Evolutionarily conserved role of oxytocin in social fear contagion in zebrafish. Science 379, 1232–1237 (2023).

26. Son, S. et al. Whole-Brain Wiring Diagram of Oxytocin System in Adult Mice. J Neurosci 42, 5021–5033 (2022).

27. Wang, P. et al. Neural Functions of Hypothalamic Oxytocin and its Regulation. ASN Neuro 14, 17590914221100706 (2022).

28. Parmaksiz, D. & Kim, Y. Navigating Central Oxytocin Transport: Known Realms and Uncharted Territories. Neuroscientist 31, 234–261 (2025).

29. Jurek, B. & Neumann, I. D. The Oxytocin Receptor: From Intracellular Signaling to Behavior. Physiological Reviews 98, 1805–1908 (2018).

30. Kang, S. J. et al. A central alarm system that gates multi-sensory innate threat cues to the amygdala. Cell Reports 40, 111222 (2022).

31. Palmiter, R. D. The parabrachial nucleus: CGRP neurons function as a general alarm. Trends Neurosci 41, 280–293 (2018).

32. Bowen, A. J. et al. Dissociable control of unconditioned responses and associative fear learning by parabrachial CGRP neurons. eLife 9, e59799 (2020).

33. Campos, C. A., Bowen, A. J., Roman, C. W. & Palmiter, R. D. Encoding of danger by parabrachial CGRP neurons. Nature 555, 617–622 (2018).

34. Palmiter, R. D. Parabrachial neurons promote nociplastic pain. Trends Neurosci 47, 722–735 (2024).

35. Condon, L. F. et al. Parabrachial *Calca* neurons drive nociplasticity. Cell Reports 43, 114057 (2024).

36. Yamada, Y., Nomura, K., Suematsu, N. & Taruno, A. Hindbrain neurons that underlie water discrimination and consumption. Curr Biol 36, 1787–1799.e6 (2026).

37. Chiang, M. C. et al. Parabrachial Complex: A Hub for Pain and Aversion. J Neurosci 39, 8225–8230 (2019).

38. Smith, M. L., Asada, N. & Malenka, R. C. Anterior cingulate inputs to nucleus accumbens control the social transfer of pain and analgesia. Science 371, 153–159 (2021).

39. Rein, B. et al. Protocols for the social transfer of pain and analgesia in mice. STAR Protoc 3, 101756 (2022).

40. Han, Y. et al. Midbrain glutamatergic circuit mechanism of resilience to socially transferred allodynia in male mice. Nat Commun 15, 4947 (2024).

41. Langford, D. J. et al. Social modulation of pain as evidence for empathy in mice. Science 312, 1967–1970 (2006).

42. Feng, C. et al. Social hierarchy modulates neural responses of empathy for pain. Soc Cogn Affect Neurosci 11, 485–495 (2016).

43. Jeon, D. et al. Observational fear learning involves affective pain system and Cav1.2 Ca2+ channels in ACC. Nat Neurosci 13, 482–488 (2010).

44. Keum, S. & Shin, H.-S. Neural Basis of Observational Fear Learning: A Potential Model of Affective Empathy. Neuron 104, 78–86 (2019).

45. Ito, W. & Morozov, A. Prefrontal-amygdala plasticity enabled by observational fear. Neuropsychopharmacology 44, 1778–1787 (2019).

46. Allsop, S. A. et al. Corticoamygdala Transfer of Socially Derived Information Gates Observational Learning. Cell 173, 1329–1342.e18 (2018).

47. Andraka, K. et al. Distinct circuits in rat central amygdala for defensive behaviors evoked by socially signaled imminent versus remote danger. Curr Biol 31, 2347–2358.e6 (2021).

48. Hernandez-Lallement, J., Gómez-Sotres, P. & Carrillo, M. Towards a unified theory of emotional contagion in rodents—A meta-analysis. Neuroscience & Biobehavioral Reviews 132, 1229–1248 (2022).

49. Puœcian, A. et al. Ability to share emotions of others as a foundation of social learning. Neuroscience & Biobehavioral Reviews 132, 23–36 (2022).

50. Oliveira, R. F. & Faustino, A. I. Social information use in threat perception: Social buffering, contagion and facilitation of alarm responses. Communicative & Integrative Biology 10, e1325049 (2017).

51. Lindström, B., Haaker, J. & Olsson, A. A common neural network differentially mediates direct and social fear learning. Neuroimage 167, 121–129 (2018).

52. Kondrakiewicz, K. et al. Social Transfer of Fear in Rodents. Current Protocols in Neuroscience 90, e85 (2019).

53. Spence, S. H. & Rapee, R. M. The etiology of social anxiety disorder: An evidence-based model. Behaviour Research and Therapy 86, 50–67 (2016).

54. Fernández, M., Mollinedo-Gajate, I. & Peñagarikano, O. Neural Circuits for Social Cognition: Implications for Autism. Neuroscience 370, 148–162 (2018).

55. Burgos-Robles, A., Gothard, K. M., Monfils, M. H., Morozov, A. & Vicentic, A. Conserved features of anterior cingulate networks support observational learning across species. Neurosci Biobehav Rev 107, 215–228 (2019).

56. Carrillo, M. et al. Emotional Mirror Neurons in the Rat’s Anterior Cingulate Cortex. Curr Biol 29, 1301–1312.e6 (2019).

57. Zhang, M.-M. et al. Glutamatergic synapses from the insular cortex to the basolateral amygdala encode observational pain. Neuron https://doi.org/10.1016/j.neuron.2022.03.030 (2022) doi:10.1016/j.neuron.2022.03.030.

58. Lischke, A. et al. Oxytocin increases amygdala reactivity to threatening scenes in females. Psychoneuroendocrinology 37, 1431–1438 (2012).

59. Zhang, X. et al. CD38-mediated oxytocin signaling in paraventricular nucleus contributes to empathic pain. Neuropharmacology 267, 110301 (2025).

60. Pauli, J. L. et al. Molecular and anatomical characterization of parabrachial neurons and their axonal projections. eLife 11, e81868 (2022).

61. Pennington, Z. T. et al. ezTrack: An open-source video analysis pipeline for the investigation of animal behavior. Sci Rep 9, 19979 (2019).

62. Demartini, C., Greco, R., Francavilla, M., Zanaboni, A. M. & Tassorelli, C. Modelling migraine-related features in the nitroglycerin animal model: Trigeminal hyperalgesia is associated with affective status and motor behavior. Physiology & Behavior 256, 113956 (2022).

63. Sureda-Gibert, P., Romero-Reyes, M. & Akerman, S. Nitroglycerin as a model of migraine: Clinical and preclinical review. Neurobiol Pain 12, 100105 (2022).

64. Barik, A., Thompson, J. H., Seltzer, M., Ghitani, N. & Chesler, A. T. A Brainstem-Spinal Circuit Controlling Nocifensive Behavior. Neuron 100, 1491–1503.e3 (2018).

65. Jaramillo, J. C. M., Aitken, C. M., Lawrence, A. J. & Ryan, P. J. Oxytocin-receptor-expressing neurons in the lateral parabrachial nucleus activate widespread brain regions predominantly involved in fluid satiation. J Chem Neuroanat 137, 102403 (2024).

66. Terburg, D. et al. The Basolateral Amygdala Is Essential for Rapid Escape: A Human and Rodent Study. Cell 175, 723–735.e16 (2018).

67. Xu, L. et al. A bottom-up septal inhibitory circuit mediates anticipatory control of drinking. Nat Neurosci 28, 2273–2284 (2025).

68. Nersesyan, Y. et al. Oxytocin modulates nociception as an agonist of pain-sensing TRPV1. Cell Rep 21, 1681–1691 (2017).

69. Yang, J. et al. Central oxytocin enhances antinociception in the rat. Peptides 28, 1113– 1119 (2007).

70. Saito, H. et al. Effects of oxytocin on responses to nociceptive and non-nociceptive stimulation in the upper central nervous system. Biochemical and Biophysical Research Communications 574, 8–13 (2021).

71. Ren, S. et al. The nociceptive inputs of the paraventricular hypothalamic nucleus in formalin stimulated mice. Neurosci Lett 841, 137948 (2024).

72. Gamal-Eltrabily, M. et al. The Rostral Agranular Insular Cortex, a New Site of Oxytocin to Induce Antinociception. J. Neurosci. 40, 5669–5680 (2020).

73. Nishimura, H. et al. Endogenous oxytocin exerts anti-nociceptive and anti-inflammatory effects in rats. Commun Biol 5, 907 (2022).

74. Li, Y.-C. et al. Distinct circuits and molecular targets of the paraventricular hypothalamus decode visceral and somatic pain. Neuron 112, 3734–3749.e5 (2024).

75. Herpertz, S. C. et al. Oxytocin Effects on Pain Perception and Pain Anticipation. The Journal of Pain 20, 1187–1198 (2019).

76. Chiang, M. C. et al. Divergent Neural Pathways Emanating from the Lateral Parabrachial Nucleus Mediate Distinct Components of the Pain Response. Neuron 106, 927–939.e5 (2020).

77. Nomura, H., Teshirogi, C., Nakayama, D., Minami, M. & Ikegaya, Y. Prior observation of fear learning enhances subsequent self-experienced fear learning with an overlapping neuronal ensemble in the dorsal hippocampus. Mol Brain 12, 21 (2019).

78. Zhang, M., Wu, Y. E., Jiang, M. & Hong, W. Cortical regulation of helping behaviour towards others in pain. Nature 626, 136–144 (2024).

79. Wolf, D. et al. Oxytocin induces the formation of distinctive cortical representations and cognitions biased toward familiar mice. Nat Commun 15, 6274 (2024).

80. Martin, L. J. et al. Reducing Social Stress Elicits Emotional Contagion of Pain in Mouse and Human Strangers. Current Biology 25, 326–332 (2015).

81. Shi, W., Fu, Y., Shi, T. & Zhou, W. Different Synaptic Plasticity After Physiological and Psychological Stress in the Anterior Insular Cortex in an Observational Fear Mouse Model. Front Synaptic Neurosci 14, 851015 (2022).

82. Li, Z. et al. Social interaction with a cagemate in pain facilitates subsequent spinal nociception via activation of the medial prefrontal cortex in rats. Pain 155, 1253–1261 (2014).

83. Mogil, J. S. Social modulation of and by pain in humans and rodents. Pain 156 **Suppl 1**, S35–S41 (2015).

84. Palagi, E., Celeghin, A., Tamietto, M., Winkielman, P. & Norscia, I. The neuroethology of spontaneous mimicry and emotional contagion in human and non-human animals. Neurosci Biobehav Rev 111, 149–165 (2020).

85. Loggia, M. L., Mogil, J. S. & Bushnell, M. C. Empathy hurts: compassion for another increases both sensory and affective components of pain perception. Pain 136, 168–176 (2008).

86. Busnelli, M., Bulgheroni, E., Manning, M., Kleinau, G. & Chini, B. Selective and Potent Agonists and Antagonists for Investigating the Role of Mouse Oxytocin Receptors. J Pharmacol Exp Ther 346, 318–327 (2013).

87. Jaramillo, A. A., Williford, K. M., Marshall, C., Winder, D. G. & Centanni, S. W. BNST transient activity associates with approach behavior in a stressful environment and is modulated by the parabrachial nucleus. Neurobiology of Stress 13, 100247 (2020).

88. Ito, M. et al. The parabrachial-to-amygdala pathway provides aversive information to induce avoidance behavior in mice. Molecular Brain 14, 94 (2021).

89. Luskin, A. T. et al. Extended amygdala-parabrachial circuits alter threat assessment and regulate feeding. Sci Adv 7, eabd3666 (2021).

90. Arthurs, J. W., Pauli, J. L. & Palmiter, R. D. Activation of Parabrachial Tachykinin 1 Neurons Counteracts Some Behaviors Mediated by Parabrachial Calcitonin Gene-related Peptide Neurons. Neuroscience 517, 105–116 (2023).

91. Barik, A. et al. A spinoparabrachial circuit defined by Tacr1 expression drives pain. Elife 10, e61135 (2021).

92. Bankhead, P. et al. QuPath: Open source software for digital pathology image analysis. Sci Rep 7, 16878 (2017).

93. Zhou, X. et al. Development of a genetically encoded sensor for probing endogenous nociceptin opioid peptide release. Nat Commun 15, 5353 (2024).

94. Barker, D. J. et al. Lateral Preoptic Control of the Lateral Habenula through Convergent Glutamate and GABA Transmission. Cell Reports 21, 1757–1769 (2017).

